# A dual-layer computational framework for prioritising therapeutic candidates targeting extracellular vesicle-mediated immune escape in pancreatic ductal adenocarcinoma

**DOI:** 10.64898/2026.07.23.740294

**Authors:** Yaojun Zhu, Xiaozhou Yang, Murtala Bindawa Isah, Xiaoying Zhang

**Author notes:** These authors contributed equally to this work (Yaojun Zhu, Xiaozhou Yang). **Corresponding author:** Dr. Xiaoying Zhang.

## Abstract

Pancreatic ductal adenocarcinoma (PDAC) is an aggressive malignancy characterised by a highly immunosuppressive tumour microenvironment and limited therapeutic responses. Tumour-derived extracellular vesicles (EVs) contribute to PDAC progression by transferring immunomodulatory molecules and tumour-associated signals, suggesting EV-associated processes as potential intervention opportunities. However, the heterogeneity of EV biology and the complexity of tumour–immune interactions make single-target intervention strategies challenging. Here, we developed a computation-driven dual-layer candidate-prioritisation framework to identify potential modulators associated with PDAC EV-mediated immune escape through complementary production-side and action-side strategies. For the production-side layer, we focused on upstream processes related to EV biogenesis, cargo regulation, inflammatory signalling, and tumour-associated pathways. An 88-gene PDAC EV-associated target framework was integrated with cell-type-resolved prognosis annotations from ctPANDA and predicted targets of 18 natural products derived from *Scutellaria baicalensis*, *Epimedium* spp., and *Cornus officinalis* to prioritise natural-product candidates with disease relevance and potential chemical tractability. In parallel, key targets with experimentally resolved ligand-binding structures were subjected to pocket-guided *de novo* small-molecule design based on co-crystal ligand-defined binding sites, followed by structural, docking-based, and physicochemical screening of generated compounds. For the action-side layer, VHH and scFv binders were computationally designed and screened against extracellular regions of MET and CD81 to prioritise candidates potentially suitable for EV recognition and capture. This study provides a computational strategy for narrowing candidate spaces across both EV-associated production pathways and released vesicle recognition. The resulting small molecules, antibody-like binder models, and screening workflows provide a resource for future experimental validation of strategies targeting PDAC EV-associated immune regulation.

**Graphical Abstract:** 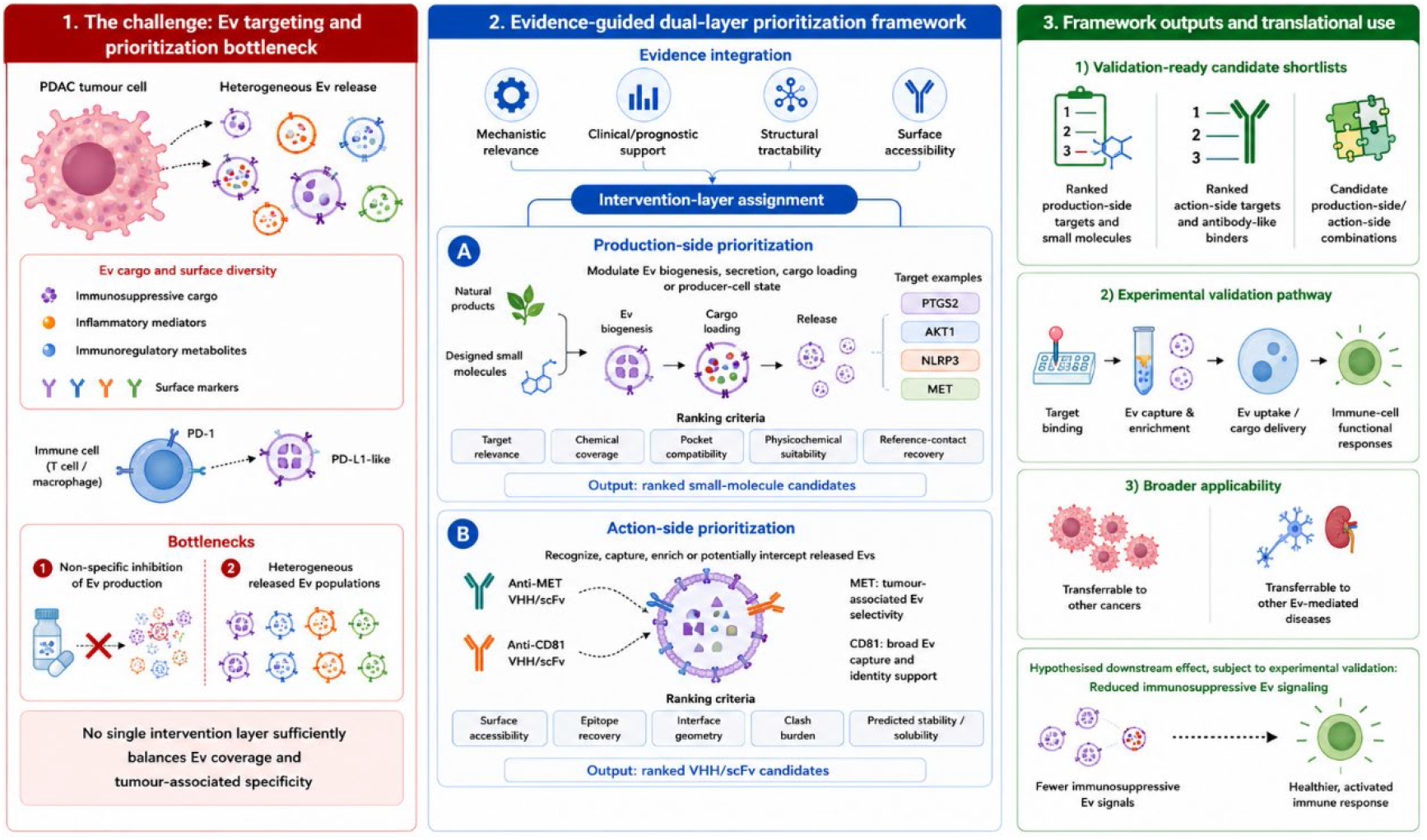

## 1 Introduction

Pancreatic ductal adenocarcinoma (PDAC) is among the gastrointestinal malignancies with the poorest prognosis [1]. Its progression is driven not only by tumour-intrinsic genetic alterations and proliferative capacity, but also by the highly complex tumour microenvironment (TME) [2]. Within this environment, tumour-derived extracellular vesicles (EVs) serve as important mediators of intercellular communication by transporting diverse biological information between cells [3, 4]. Through the transfer of proteins, lipids, miRNAs, cytokines, immunomodulatory molecules, and other bioactive cargoes, EVs can reshape the TME and contribute to immune escape, metastatic progression, and therapeutic resistance [5, 6]. EVs are therefore increasingly recognised as functional mediators of PDAC progression and as potential subjects for biomarker discovery and therapeutic investigation.

However, most current EV-targeting strategies focus on only one step of EV-mediated communication. Blocking EV production or release can reduce the number of new vesicles, but it cannot remove the pro-tumour EVs that are already present in the tumour microenvironment or circulation. EV secretion also depends on several overlapping pathways. When one pathway is blocked, tumour cells may continue to release active EVs through other routes or shift to compensatory secretion mechanisms [7, 8]. The opposite approach has a similar limitation. Blocking EV recognition or uptake may reduce their effects on recipient cells, but it does not stop tumour cells from producing more immunosuppressive EVs. These vesicles can continue to accumulate and replenish the extracellular pool, which may weaken the depth and duration of the intervention [9, 10].

To address these challenges, this study proposes a dual-layer candidate-prioritisation framework that explores complementary strategies targeting different stages of PDAC EV-associated immune regulation. For the production-side layer, natural-product-derived compounds are screened to identify candidates potentially associated with tumour-cell pathways involved in EV biogenesis, secretion, cargo regulation, inflammatory signalling, and tumour-related processes. In parallel, target-guided small-molecule candidates are designed using experimentally resolved protein structures and ligand-defined binding pockets to explore additional chemical candidates for downstream evaluation. For the action-side layer, VHH and scFv binders are computationally designed against extracellular regions of MET and CD81 to prioritise potential candidates for future EV recognition, enrichment, or functional interception studies. This framework aims to narrow the candidate space for experimental validation by integrating disease-associated target information, small-molecule prioritisation, and antibody-like binder design. This framework provides a computational foundation for future research on multi-stage strategies targeting PDAC EV-related immune regulation.

## 2 Materials and methods

### 2.1 Construction of the PDAC EV-associated candidate target framework

A literature search was performed in Web of Science (up to June 12, 2026) using the following keywords: “pancreatic ductal adenocarcinoma,” “extracellular vesicle,” “exosome,” “immune escape,” “cargo loading,” “biogenesis,” and “surface marker.” Candidate genes were included based on published experimental evidence supporting their relevance to PDAC biology, EV-associated functions, or related tumour-microenvironmental processes. A manually curated PDAC EV-associated candidate target framework was assembled from previously reported studies [11–24]. Each retained gene was annotated using two complementary classification systems. First, genes were assigned to one of eight functional modules according to their reported biological roles: EV biogenesis and release, cargo loading, immunosuppressive cargo, PDAC-associated surface markers, immune checkpoint and T-cell dysfunction, myeloid and macrophage regulation, inflammatory and oncogenic signalling, and extracellular matrix and stromal remodelling. Second, genes were categorized into production-side or action-side layers according to their potential roles within the candidate-prioritisation framework. The production-side layer included genes potentially associated with EV generation, secretion, cargo regulation, or the immunoregulatory state of EV-producing cells. The action-side layer included EV-associated or surface-accessible candidates potentially suitable for post-release EV recognition, enrichment, or interception studies.

### 2.2 Cell-type-resolved prognostic annotation

The curated PDAC EV-associated candidate target framework was integrated with cell-type-resolved prognosis data from ctPANDA [25]. The ctPANDA dataset was processed into a standardised table containing gene symbols, cell types, and prognosis annotations. Only entries labelled as “Worse” or “Better” were retained, while records without prognosis classification were excluded. Gene matching was performed using official gene symbols. For each matched gene, prognosis associations across different cell types were summarized and classified as Worse-only, Better-only, or mixed. Worse-only and Better-only categories represented consistent prognosis directions across retained cell-type annotations, whereas mixed classification indicated opposite prognosis associations among different cell types.

### 2.3 Natural-product library curation and target prediction

A natural-product compound library was assembled from *Scutellaria baicalensis*, *Epimedium* spp., and *Cornus officinalis*. The library included representative flavones, flavonol glycosides, iridoid glycosides, and triterpenoids. For each compound, PubChem identifiers, molecular formulae, molecular weights, canonical SMILES, InChIKeys, and structural files were collected. Canonical SMILES were submitted to SwissTargetPrediction [26] using the *Homo sapiens* prediction setting. The resulting compound–target prediction records were merged and standardised according to gene symbols, UniProt identifiers, ChEMBL identifiers, target classes, and prediction probability values. Records containing missing, zero, or non-numeric probability values, or lacking valid target annotations, were excluded. Predicted compound targets were intersected with the PDAC EV-associated candidate target framework using official gene symbols. Compound–target associations were summarized according to the number of associated compounds, contributing plant sources, maximum prediction probability, and mean prediction probability.

### 2.4 Protein association network and pathway enrichment analysis

Genes shared between the ctPANDA-supported candidate framework and natural-product target prediction were submitted to STRING [27] for protein association network analysis. The organism was restricted to *Homo sapiens*, and the minimum interaction confidence score was set to 0.400. Network information, including nodes, edges, and degree values, was exported for visualisation. Functional enrichment analysis was performed using the STRING enrichment module. Gene Ontology biological process, KEGG pathway, and Reactome pathway terms with a false discovery rate (FDR) < 0.05 were considered significant and retained for visualisation.

### 2.5 Receptor and ligand preparation

PTGS2, MMP2, NLRP3, MET, and AKT1 were selected for molecular docking because they satisfied three criteria: (i) inclusion in the prioritised PDAC EV-associated candidate target framework, (ii) predicted coverage by the natural-product compound library, and (iii) availability of experimentally resolved ligand-bound protein structures. The receptor structures used for docking were PTGS2–COH (PDB ID: 5IKR), MMP2–B3P (PDB ID: 7XJO), NLRP3–RM5 (PDB ID: 7ALV), MET–DF6 (PDB ID: 3VW8), and AKT1–XM1 (PDB ID: 3OCB). Protein structures were prepared by removing non-essential water molecules, solvents, buffers, and unrelated heteroatoms. Cofactors and metal ions required for maintaining ligand-binding pocket geometry were retained. Protein structures were converted into PDBQT format using Meeko [28]. Ligand structures were generated in three-dimensional conformations, hydrogen-completed, and protonated at pH 7.4 before conversion into PDBQT format.

### 2.6 Molecular docking

Molecular docking was performed using AutoDock Vina v1.2.7 [29]. Each selected natural product was docked into the corresponding co-crystal ligand-defined binding pocket. Docking calculations were performed with an exhaustiveness value of 32 and 10 output modes. The mode-1 Vina score was extracted for within-target ranking. Vina scores were interpreted as relative docking scores obtained under the same target-specific docking protocol rather than direct measurements of binding affinity. Protein residues within 4 Å of the docked ligand were defined as ligand-contact residues. The cognate ligand from each crystal structure was independently redocked using the same receptor preparation, docking box definition, and Vina parameters. Heavy-atom RMSD values between redocked and experimentally observed ligand poses were calculated to evaluate docking-pose recovery.

### 2.7 Pocket-guided *de novo* small-molecule generation and screening

Pocket-guided small-molecule generation was performed using AnewOmni [30] based on ligand-defined binding pockets derived from NLRP3–RM5, AKT1–XM1, MET–DF6, and PTGS2–ID8 complexes. The PTGS2–ID8 system was used exclusively for molecular generation and redocking and was analysed separately from the PTGS2–COH natural-product docking workflow. Generated molecules were initially evaluated using RDKit[33] for molecular validity, three-dimensional coordinate completeness, molecular weight, LogP, topological polar surface area (tPSA), hydrogen-bond donor count, hydrogen-bond acceptor count, rotatable-bond count, quantitative estimate of drug-likeness (QED), and pan-assay interference structure (PAINS) alerts.

Protein–ligand complexes were further evaluated using geometric criteria, including minimum protein–ligand heavy-atom distance, contact-residue number, ligand-contact atom fraction, and severe steric clashes. Retained candidates were redocked into the corresponding receptor pocket using the reference ligand-defined docking box. Reference-pocket similarity was evaluated using reference-contact recovery and Jaccard similarity based on ligand-contact residue profiles. Candidates were classified into report-priority, physicochemical-review, docking-only-review, or not-selected categories according to predefined structural, physicochemical, clash, contact-overlap, and docking criteria.

### 2.8 Antibody-like binder design and computational screening

VHH and scFv candidates were designed using Rfantibody [14]. MET-directed binder design used anti-MET structural templates from PDB IDs 5LSP and 6I04. CD81-directed binder design used the CD81–5A6 Fab complex (PDB ID: 6U9S) and CD81–K13 scFv complex (PDB ID: 5DFW). MET was selected as a PDAC-associated surface marker candidate, whereas CD81 was selected as an EV-associated membrane marker frequently used for EV recognition studies. Antigen chains were standardised as chain R. Binder chains were renamed as H/L for Fab-derived templates or H for single-domain binder templates. Interface hotspots were defined as antigen residues located within 4 Å of reference binder atoms. When more than 15 antigen-contact residues were identified, the 15 nearest residues were retained as design hotspots. Candidate sequences were generated using RFantibody and ProteinMPNN [31], followed by structural evaluation using RoseTTAFold2 [32]. Candidate binders were screened according to epitope recovery, complex interface predicted TM-score (iPTM), interface predicted aligned error (ipAE), hotspot recovery, severe steric clashes, interface RMSD, PRODIGY-derived interaction estimates [33], FoldX-derived stability estimates [34], ProtParam-derived N-end-rule half-life prediction [35], and Protein-Sol-derived sequence-based solubility scores [36].

## 3 Results

### 3.1 Construction and cell-type-resolved prognostic annotation of the PDAC EV-associated candidate target framework

A total of 88 genes associated with PDAC EV biology were curated and organised into eight functional modules (Table 1). These modules covered EV biogenesis and release, cargo loading, immunosuppressive cargo, tumour-associated surface markers, immune dysfunction, myeloid regulation, inflammatory and oncogenic signalling, and stromal remodelling. Together, they represented the major stages through which EVs may contribute to immune escape, from vesicle production and cargo incorporation to interactions with recipient cells and the tumour microenvironment.

**Table 1.**
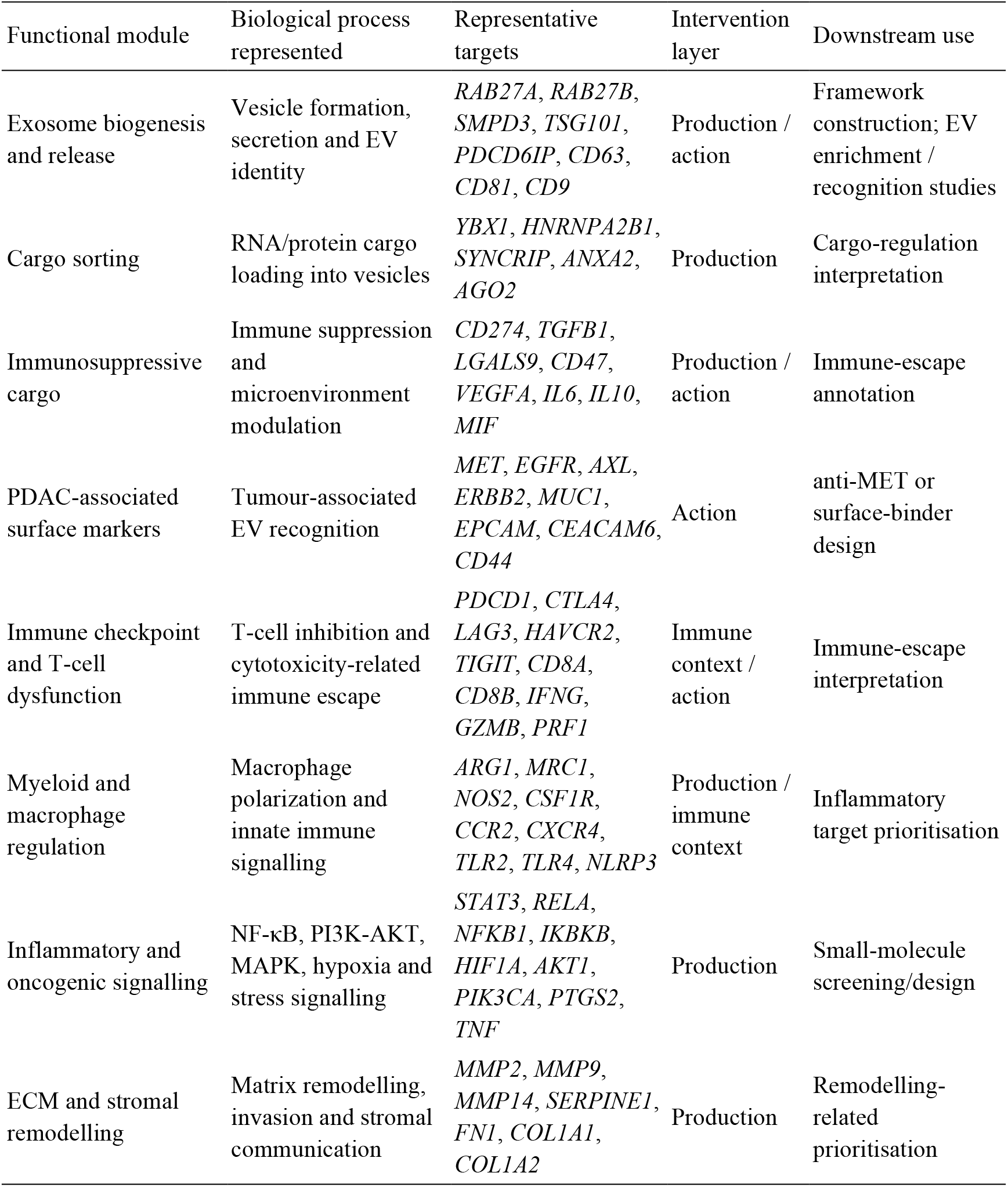
Curation of an 88-gene PDAC EV-associated immune-regulation candidate target framework.

Integration of the 88-gene framework with ctPANDA cell-type-resolved prognosis data identified 42 matched genes and 81 gene–cell-type prognosis records (Table 2). Among the matched genes, 24 were associated exclusively with worse prognosis, 10 showed opposite prognostic directions across different cell types, and 8 were associated exclusively with better prognosis. PLOD2 showed the broadest adverse pattern, with worse-prognosis associations in eight cell types. Several EV-related and signalling genes showed mixed associations, indicating that their prognostic relevance varied across cellular compartments.

**Table 2.** Representative ctPANDA-prioritised targets in the PDAC EV-associated immune-regulation framework.

| Target | Functional module | Intervention layer | Clinical pattern | Suggested interpretation |
| --- | --- | --- | --- | --- |
| *PLOD2* | PDAC stroma/ECM remodelling | Production-side / stromal mechanism | Worse-only | Cell-type-resolved prognosis-associated stromal-remodelling candidate |
| *SERPINE1* | PDAC stroma/ECM remodelling | Production-side / stromal mechanism | Worse-only | Production-side remodelling-associated candidate |
| *CHMP4B* | EV biogenesis and release | Production-side | Worse-only | Clinically supported EV biogenesis and release candidate |
| *CD63* | EV biogenesis and release | Action-side / exosome marker | Worse-only | EV recognition/enrichment marker |
| *CD9* | EV biogenesis and release | Action-side / exosome marker | Worse-only | EV recognition/enrichment marker |
| *DDX3X* | Exosome cargo loading | Production-side | Worse-only | Clinically supported cargo-related candidate |
| *LOX* | PDAC stroma/ECM remodelling | Production-side / stromal mechanism | Worse-only | ECM remodelling-associated candidate |
| *TGFB1* | Immunosuppressive cargo | Production / immune context | Worse-only | Immunosuppressive exosomal cargo-related candidate |
| *VEGFA* | Immunosuppressive cargo | Production / immune context | Worse-only | Angiogenesis and immune microenvironment-related candidate |
| *AXL* | PDAC tumour/exosome surface marker | Action-side / context-dependent | Worse-only | Surface-accessible candidate for tumour-associated exosome interpretation |
| *ITGB1* | PDAC tumour/exosome surface marker | Action-side / context-dependent | Mixed | Context-dependent surface marker; requires cautious interpretation |
| *ANXA2* | Exosome cargo loading | Production-side / cargo-related | Mixed | Context-dependent cargo-loading candidate |
| *CD44* | PDAC tumour/exosome surface marker | Action-side / context-dependent | Mixed | Context-dependent tumour/exosome surface marker |
| *RELA* | Core signalling pathway | Production-side | Mixed | NF-κB-related signalling candidate requiring cell-type-aware interpretation |
| *AKT1* | Core signalling pathway | Production-side | Mixed | Intracellular signalling candidate; interpret cautiously |

For downstream analysis, 32 prognosis-associated genes were assigned to the production-side layer and 10 to the action-side layer (Table 3). Production-side genes were mainly associated with EV formation, cargo regulation, intracellular signalling, immunosuppression, and stromal remodelling. Action-side genes mainly represented surface-accessible or EV-associated molecules that could support EV recognition, capture, or blockade. Worse-prognosis records predominated in both layers. However, genes with mixed or better-prognosis associations were also retained because their effects varied according to cell type. This integrated framework provided the basis for subsequent small-molecule prioritisation and binder design.

**Table 3.**
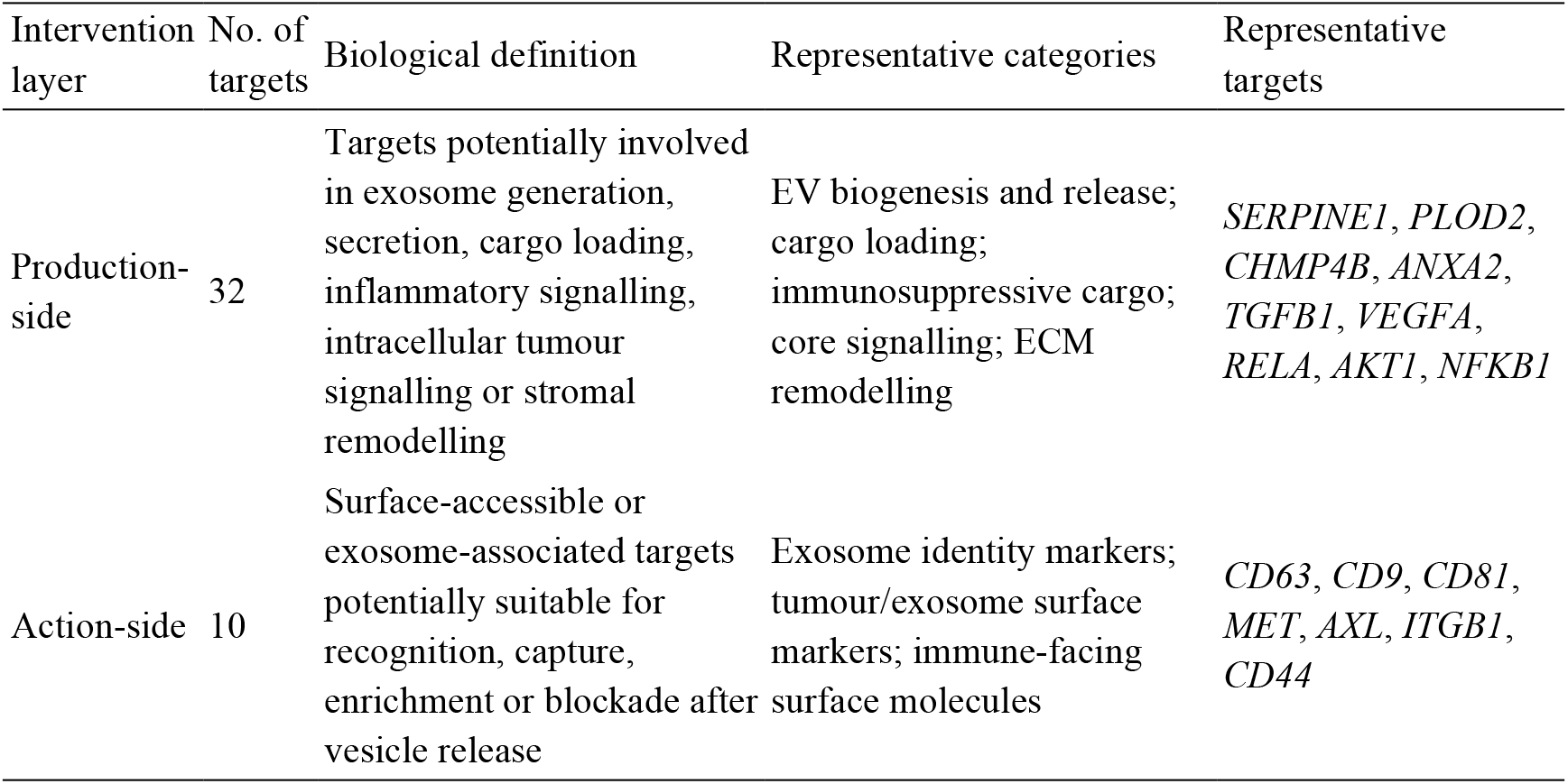
Production-side and action-side stratification of ctPANDA-supported EV-associated candidates.

### 3.2 Natural-product target prediction, protein association analysis, and molecular docking

#### 3.2.1 Natural-product target overlap and functional characterisation of prioritised candidates

The natural-product library contained 18 compounds derived from *Scutellaria baicalensis*, *Epimedium* spp., and *Cornus officinalis* (Table 4). SwissTargetPrediction generated 1,800 initial compound–target records. After data cleaning, 1,726 records involving 342 unique predicted genes were retained. Integration with the 42 ctPANDA-supported genes identified 62 compound–target associations involving 13 overlapping candidates.

**Table 4.** Natural products included in the chemical library.

| Herb source | Compound | Chemical class | PubChem CID | Molecular formula | Molecular weight (Da) |
| --- | --- | --- | --- | --- | --- |
| *Scutellaria baicalensis* | baicalein | Flavone | 5281605 | C<sub>15</sub>H<sub>10</sub>O<sub>5</sub> | 270.24 |
|  | baicalin | Flavone glycoside | 64982 | C<sub>21</sub>H<sub>18</sub>O<sub>11</sub> | 446.40 |
|  | wogonin | Flavone | 5281703 | C<sub>16</sub>H<sub>12</sub>O<sub>5</sub> | 284.26 |
|  | wogonoside | Flavone glycoside | 3084961 | C<sub>22</sub>H<sub>20</sub>O<sub>11</sub> | 460.40 |
|  | oroxylin A | Flavone | 5320315 | C<sub>16</sub>H<sub>12</sub>O<sub>5</sub> | 284.26 |
|  | scutellarein | Flavone | 5281697 | C<sub>15</sub>H<sub>10</sub>O<sub>6</sub> | 286.24 |
| *Epimedium* spp. | icariin | Prenylated flavonol glycoside | 5318997 | C<sub>33</sub>H<sub>40</sub>O<sub>15</sub> | 676.70 |
|  | icaritin | Prenylated flavonol | 5318980 | C<sub>21</sub>H<sub>20</sub>O<sub>6</sub> | 368.40 |
|  | baohuoside I | Prenylated flavonol glycoside | 5488822 | C<sub>27</sub>H<sub>30</sub>O<sub>10</sub> | 514.50 |
|  | epimedin A | Prenylated flavonol glycoside | 53486399 | C<sub>40</sub>H<sub>52</sub>O<sub>19</sub> | 836.80 |
|  | epimedin B | Prenylated flavonol glycoside | 5748393 | C<sub>38</sub>H<sub>48</sub>O<sub>19</sub> | 808.80 |
|  | epimedin C | Prenylated flavonol glycoside | 5748394 | C<sub>39</sub>H<sub>50</sub>O<sub>19</sub> | 822.80 |
| *Cornus officinalis* | loganin | Iridoid glycoside | 87691 | C<sub>17</sub>H<sub>26</sub>O<sub>10</sub> | 390.40 |
|  | morroniside | Iridoid glycoside | 11228693 | C<sub>17</sub>H<sub>26</sub>O<sub>11</sub> | 406.40 |
|  | cornuside | Iridoid glycoside | 131348 | C<sub>24</sub>H<sub>30</sub>O<sub>14</sub> | 542.50 |
|  | ursolic acid | Triterpenoid | 64945 | C<sub>30</sub>H<sub>48</sub>O<sub>3</sub> | 456.70 |
|  | oleanolic acid | Triterpenoid | 10494 | C<sub>30</sub>H<sub>48</sub>O<sub>3</sub> | 456.70 |
|  | sweroside | Iridoid glycoside | 161036 | C<sub>16</sub>H<sub>22</sub>O<sub>9</sub> | 358.34 |

Of these 13 candidates, nine were assigned to the production-side layer and four to the action-side layer. The production-side candidates included *SERPINE1*, *RELA*, *PTGS2*, *MMP2*, *AKT1*, *TNF*, *NLRP3*, *FYN*, and *TP53*, whereas the action-side candidates comprised *MET*, *CD81*, *AXL*, and *LGALS9*. Among the overlapping genes, *PTGS2* showed the broadest compound coverage and was predicted for 17 compounds. *MMP2* was covered by 11 compounds, whereas *AKT1* and *TNF* were each covered by seven compounds. *MET* showed the highest coverage among the action-side candidates, with five predicted compounds (Figure 1A). At the compound level, cornuside covered seven framework-associated genes, followed by baohuoside I with six genes.

**Figure 1.**
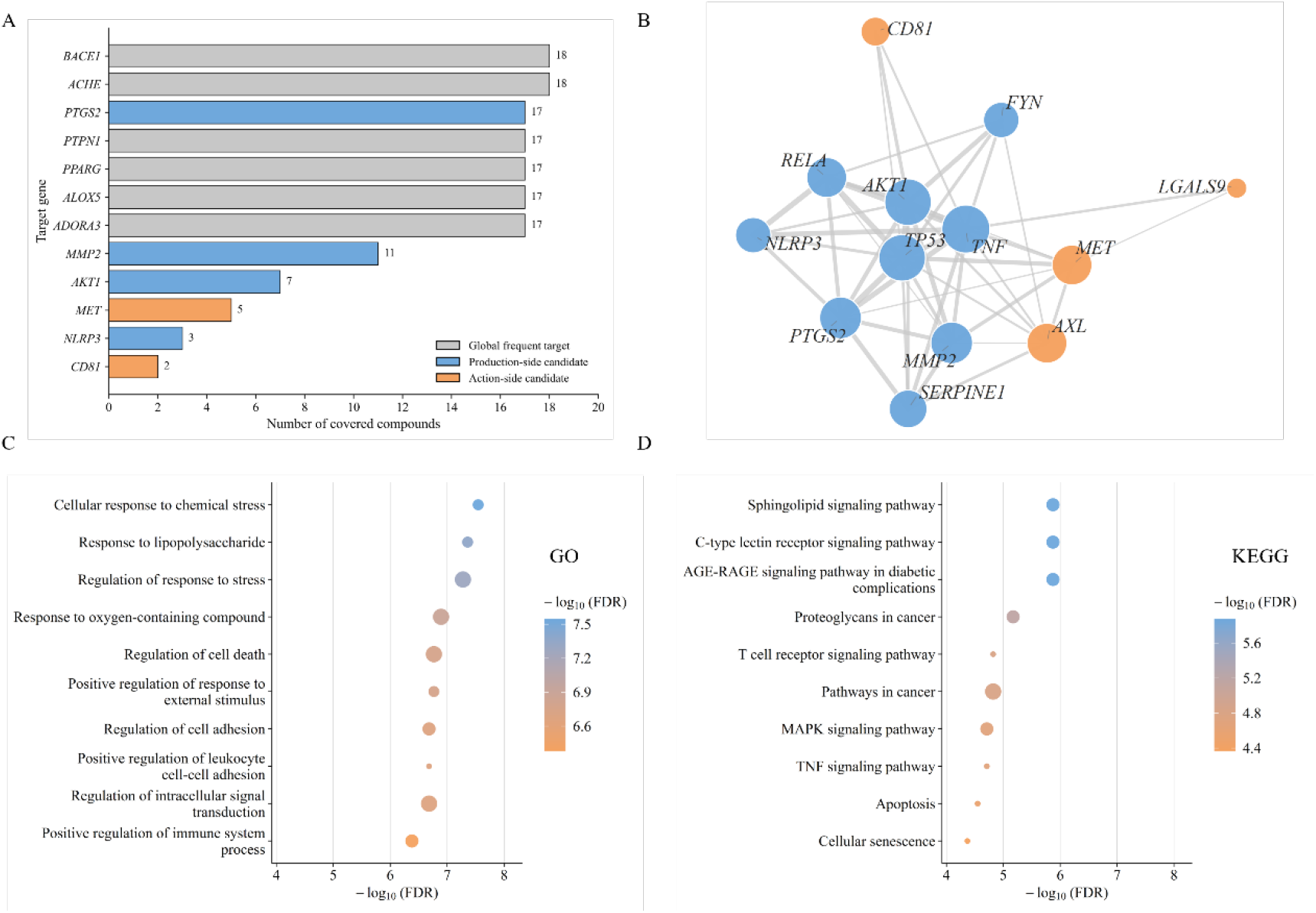
Predicted target coverage, protein association network, and enrichment analysis of natural-product-associated candidates. **Note:** (A) Compound-level coverage of predicted targets within the PDAC EV-associated candidate framework. Grey bars indicate frequently predicted targets, whereas blue and orange bars indicate production-side and action-side candidates, respectively. Values represent the number of compounds predicted to cover each target. (B) STRING protein–protein association network of the 13 overlapping candidate genes identified from natural-product target prediction and ctPANDA-supported annotation. Node colour indicates candidate category, node size represents network degree, and edge thickness represents interaction confidence. (C–D) GO biological process and KEGG pathway enrichment analyses of the overlapping target set. Point position represents −log_10_(FDR), point size indicates gene number, and colour represents enrichment significance.

The 13 overlapping candidates formed a connected STRING protein-association network containing 46 edges (Figure 1B). TNF showed the highest network degree, followed by AKT1 and TP53, indicating that inflammatory and oncogenic signalling proteins occupied central positions within the network. Functional enrichment linked the candidate set to cellular stress responses, inflammatory regulation, immune signalling, cell death, cell adhesion, and intracellular signal transduction (Figure 1C). The enriched pathways included sphingolipid signalling, C- type lectin receptor signalling, T-cell receptor signalling, MAPK signalling, and TNF signalling, together with pathways related to cancer, apoptosis, and cellular senescence (Figure 1D). Overall, the predicted natural-product targets converged on an interconnected set of production-side and action-side candidates relevant to PDAC EV-associated immune regulation, providing the basis for subsequent molecular docking analysis.

#### 3.2.2 Molecular docking and interaction profiles of representative natural product–target pairs

Five structurally resolved targets, PTGS2, MMP2, NLRP3, MET, and AKT1, were selected for molecular docking. Seven representative natural products were evaluated, generating 35 compound–target combinations. The mode-1 Vina scores ranged from −5.509 to −9.846 kcal mol^-1^ (Figure 2).

**Figure 2.**
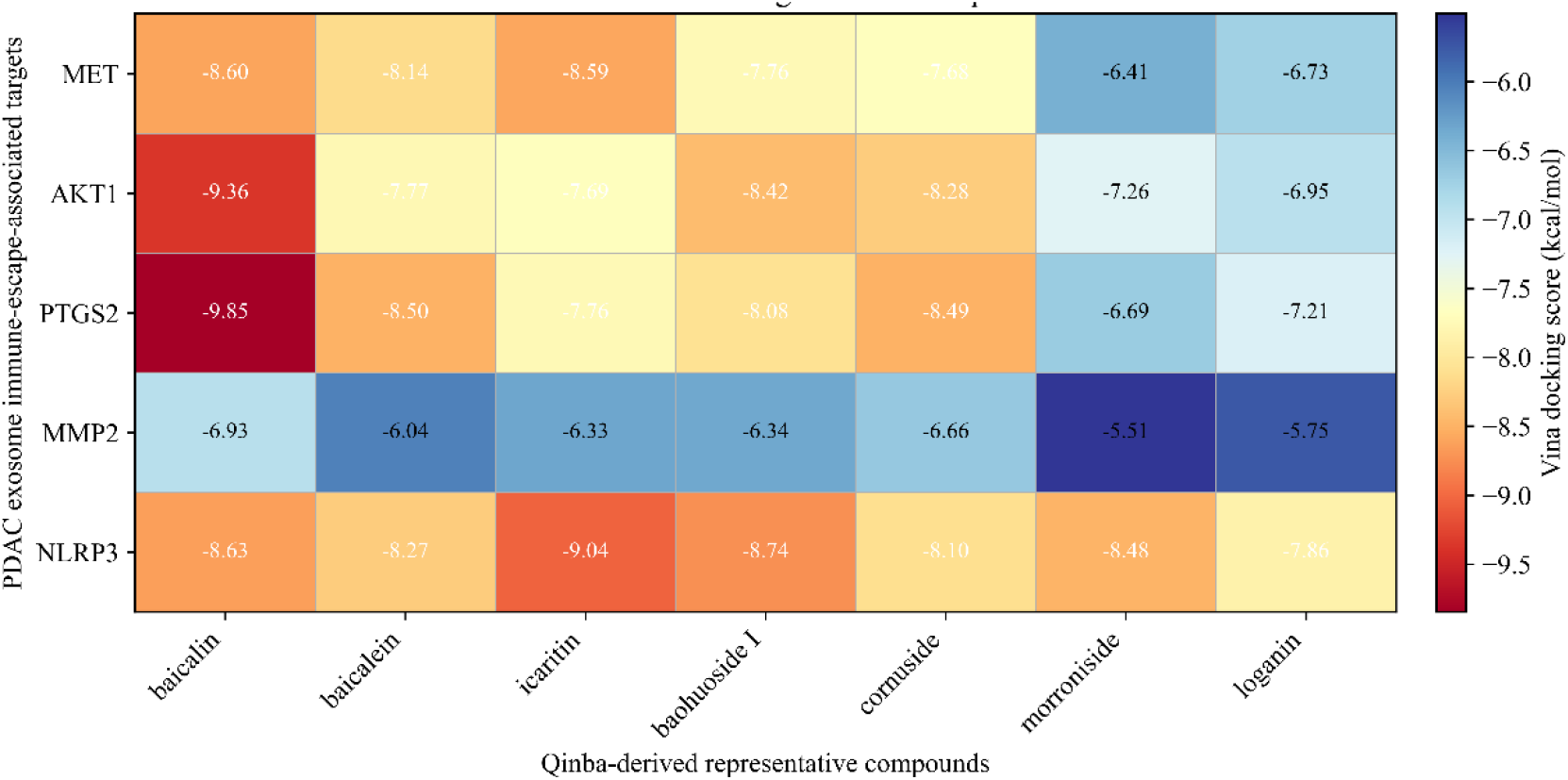
Mode-1 Vina scores of representative natural-product compounds docked to selected PDAC EV-associated candidate targets. **Note:** The heatmap shows mode-1 Vina docking scores of representative natural-product compounds against *MET*, *AKT1*, *PTGS2*, *MMP2*, and *NLRP3*. Values are shown in kcal mol^-1^. More negative values represent more favourable docking scores under the same target-specific docking protocol.

Baicalin showed the most favourable score for PTGS2, AKT1, MET, and MMP2 under the corresponding target-specific docking protocols. The strongest overall score was observed for the PTGS2–baicalin complex at −9.846 kcal mol^-1^, followed by AKT1–baicalin at −9.363 kcal mol^-1^. For NLRP3, icaritin produced the most favourable score at −9.039 kcal mol^-1^, and six of the seven tested compounds scored below −8.0 kcal mol^-1^. In contrast, docking to MMP2 was consistently weaker, with scores ranging from −5.509 to −6.930 kcal mol^-1^. These results identified PTGS2– baicalin, AKT1–baicalin, NLRP3–icaritin, and MET–baicalin as the most informative complexes for structural examination.

The representative poses occupied the predefined ligand-binding regions and formed combinations of polar, hydrophobic, and π-related contacts with surrounding residues (Figures 3–6). The PTGS2–baicalin and NLRP3–icaritin complexes displayed extensive contact networks within their respective pockets, whereas the MET–baicalin and AKT1–baicalin poses also showed geometrically compatible interactions with key pocket residues. Together, the docking scores and interaction profiles supported the structural compatibility of these natural products with the selected target pockets and provided a basis for subsequent candidate comparison and target-guided molecular design.

**Figure 3.**
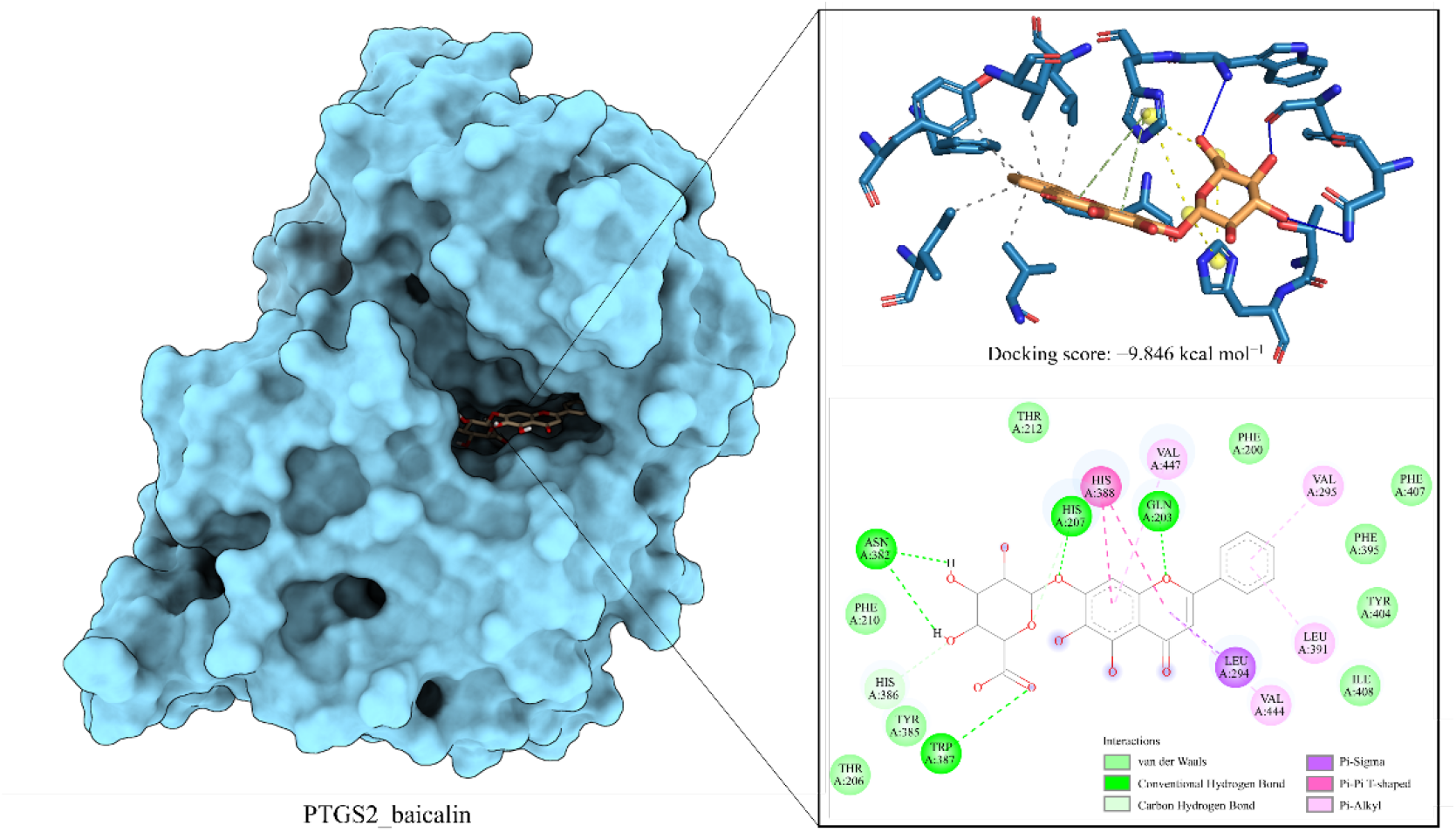
Docking pose and contact-residue analysis of the PTGS2–baicalin complex. **Note:** The left panel shows the surface representation of *PTGS2* with baicalin positioned within the ligand-defined binding pocket. The upper-right panel presents a magnified three-dimensional view of the predicted docking pose and surrounding pocket residues. The lower-right panel displays a two-dimensional receptor–ligand interaction map. The PTGS2–baicalin docking score was −9.846 kcal mol^-1^. Baicalin formed hydrogen-bond-related and hydrophobic contacts with residues including *GLN203*, *HIS207*, *ASN382*, *LEU294*, *VAL295*, *TRP387*, *HIS388*, *VAL444*, and *VAL447*. These contacts indicate geometric compatibility of the predicted pose within the PTGS2 pocket under the applied docking protocol.

**Figure 4.**
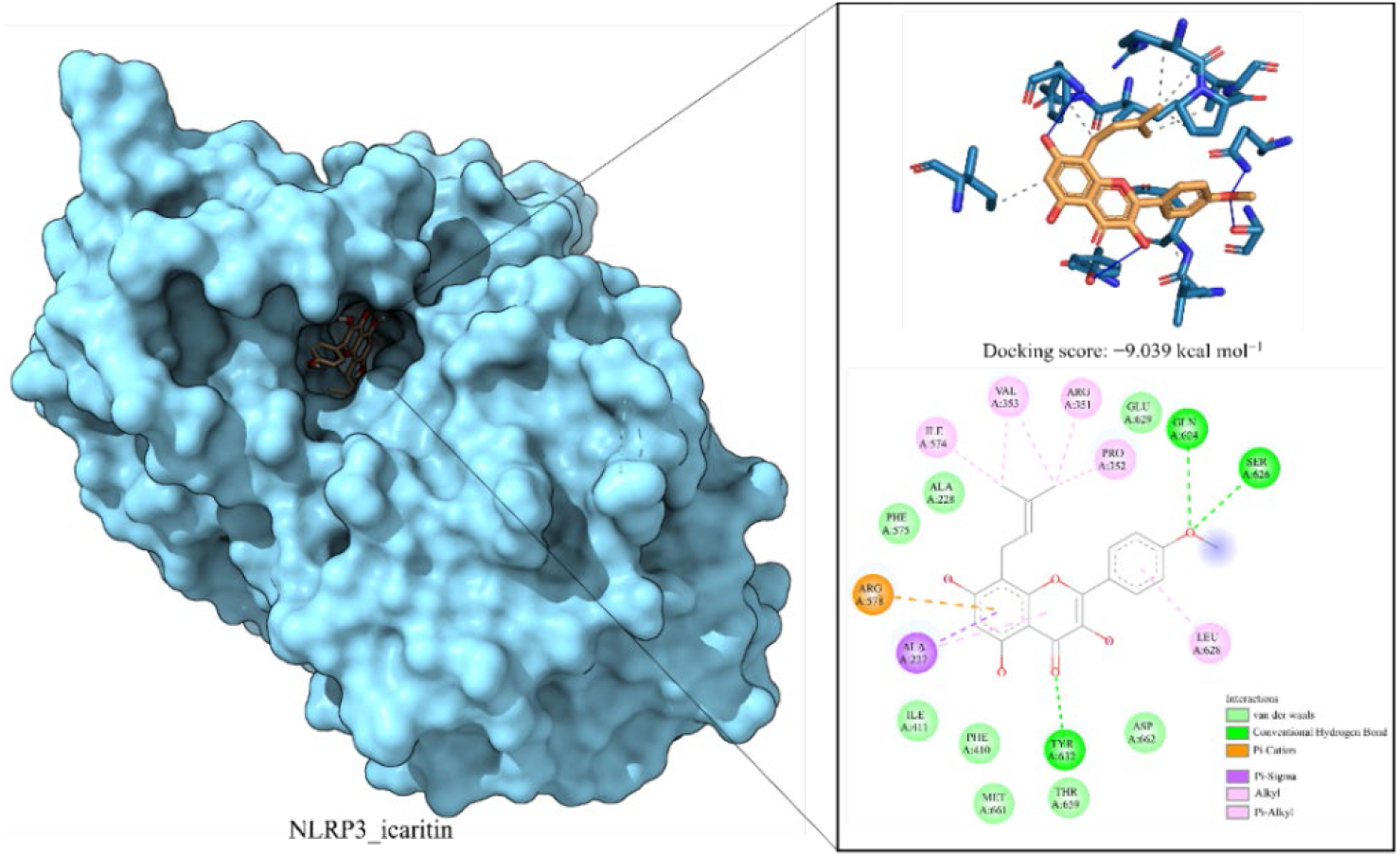
Docking pose and contact-residue analysis of the NLRP3–icaritin complex. **Note:** The left panel shows the surface representation of *NLRP3* with icaritin accommodated within the ligand-defined binding region. The upper-right panel presents the enlarged three-dimensional docking pose. The lower-right panel shows the receptor–ligand interaction map. The NLRP3–icaritin docking score was −9.039 kcal mol^-1^. Icaritin formed multiple polar and hydrophobic contacts with residues including *GLU369*, *GLN366*, *SER626*, *TYR412*, *ARG578*, *ALA227*, *ILE574*, *VAL353*, *ARG351*, *PRO352*, and *LEU628*. These interactions indicate compatibility of the predicted pose with the NLRP3 pocket under the applied docking protocol.

**Figure 5.**
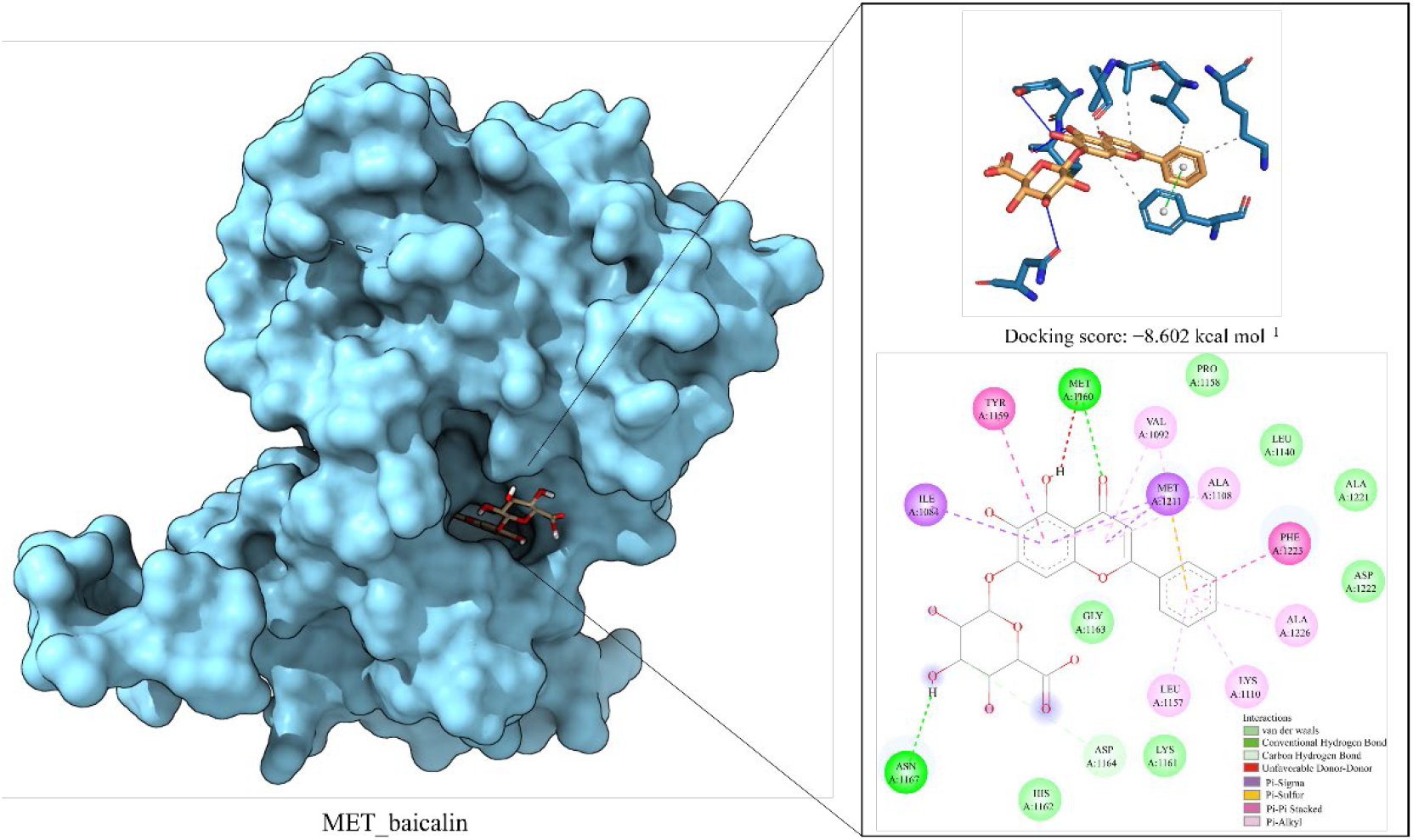
Representative docking mode and interaction profile of the baicalin–MET complex. **Note:** The left panel shows the surface representation of *MET* with baicalin positioned within the defined binding pocket. The upper-right panel displays the enlarged three-dimensional docking pose with surrounding residues. The lower-right panel presents the two-dimensional receptor–ligand interaction map. The baicalin–MET docking score was −8.602 kcal mol^-1^. Baicalin formed polar and hydrophobic contacts with residues including *ASN1168*, *MET1160*, and *GLY1163*, together with additional interactions involving *ILE1084*, *MET1121*, *VAL1092*, *ALA1108*, *PHE1123*, *ILE1157*, and *TYR1159*. These contacts support geometric compatibility of the predicted baicalin pose within the MET pocket.

**Figure 6.**
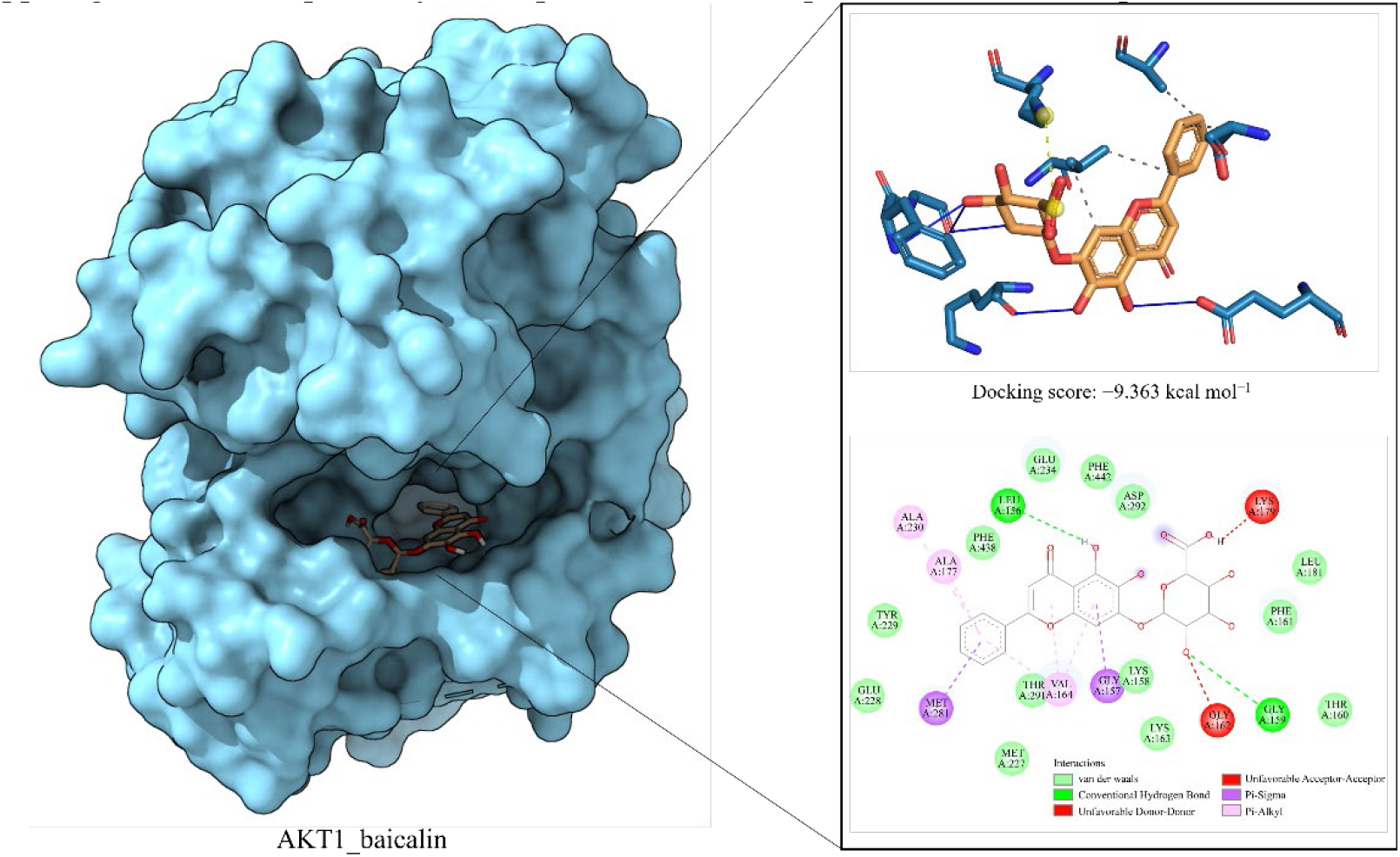
Representative docking mode and interaction profile of the baicalin–AKT1 complex. **Note:** The left panel shows the surface representation of *AKT1* with baicalin positioned within the predicted binding region. The upper-right panel provides an enlarged three-dimensional view of the docking pose and surrounding residues. The lower-right panel presents the two-dimensional receptor–ligand interaction map. The baicalin–AKT1 docking score was −9.363 kcal mol^-1^. Baicalin formed hydrogen-bond-related, π-related, and hydrophobic contacts with residues including *GLU234*, *LEU156*, *GLY159*, *PHE161*, *LEU181*, *GLY157*, *THR164*, *MET281*, *GLU228*, *TYR229*, *ALA177*, and *ALA230*. These interactions indicate compatibility of the predicted pose within the AKT1 pocket under the applied docking protocol.

### 3.3 Target-guided design of small-molecule candidates using co-crystal-defined binding pockets

#### 3.3.1 RM5-pocket-guided generation and prioritisation of NLRP3 small-molecule candidates

The experimentally resolved RM5-binding pocket of NLRP3 was used to generate 100 candidate protein–ligand complexes. Initial ligand-level and complex-level quality control retained 99 candidates, while one candidate was flagged because of a PAINS alert. Subsequent inspection of pocket occupancy, protein–ligand contacts, ligand completeness, and steric clashes reduced the set to 88 candidates for redocking. No candidate was excluded because of severe clashes, ligand displacement, or missing ligand atoms.

All 88 candidates were successfully redocked into the RM5-defined pocket using the same receptor preparation, docking box, and AutoDock Vina parameters. Their mode-1 Vina scores ranged from −10.690 to −6.012 kcal mol^-1^. Six candidates were classified as very strong, 21 as strong, 35 as good, 21 as moderate, and five as weak (Figure 7A, B). Integration of docking performance with physicochemical properties, structural quality, and pocket-contact features identified 16 report-priority candidates. A further seven candidates required physicochemical review, four were retained for docking-only review, and 61 were not selected.

**Figure 7.**
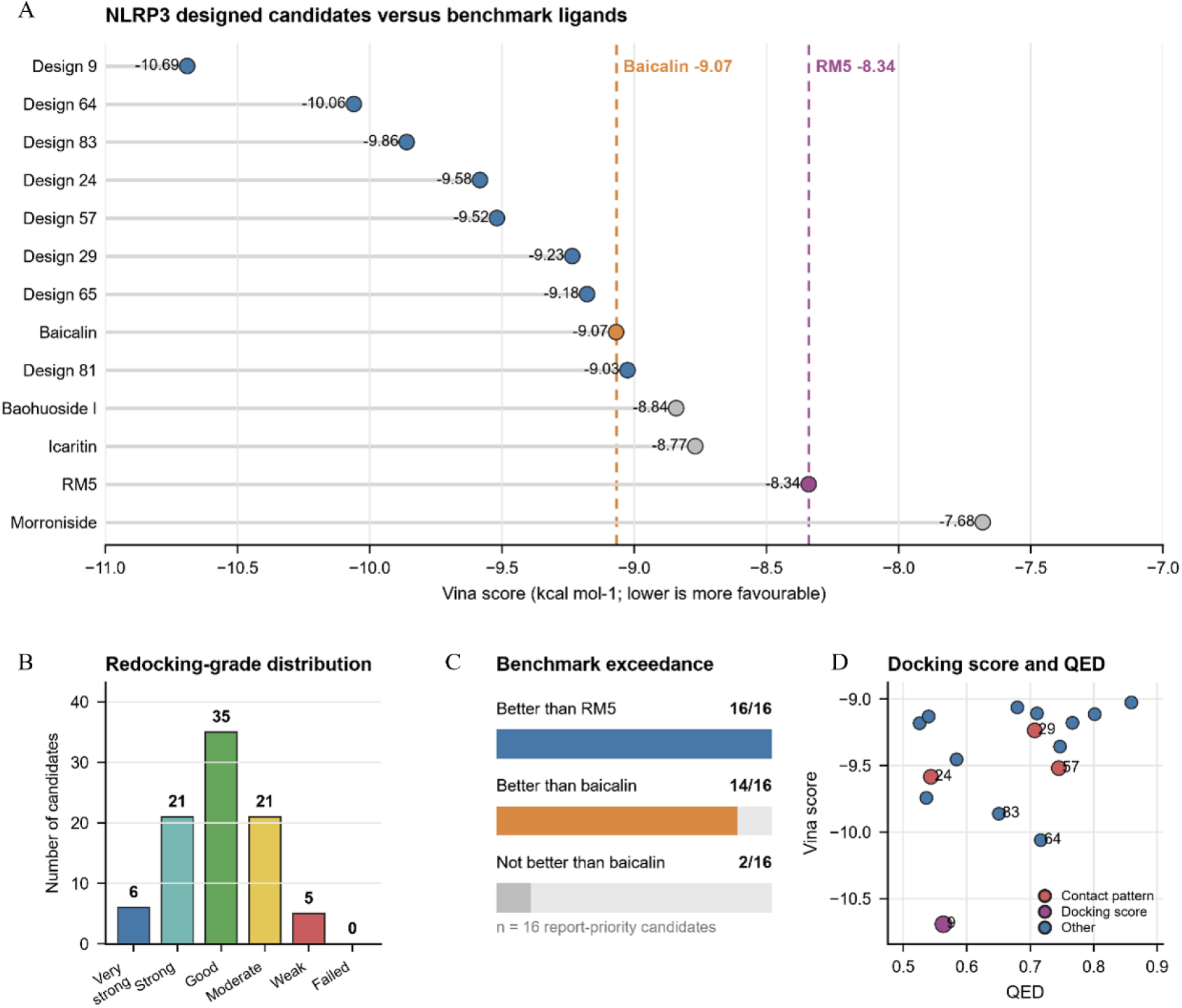
Redocking-based prioritisation of NLRP3 small-molecule candidates in the RM5-defined binding pocket. **Note:** (A) Mode-1 Vina scores of representative NLRP3-designed candidates and benchmark ligands redocked into the RM5-defined pocket. Dashed lines indicate RM5 and baicalin reference scores. (B) Distribution of redocking-score categories among the 88 final candidates. (C) Comparison of report-priority candidates with RM5 and baicalin benchmarks. (D) Relationship between QED values and Vina scores for the 16 report-priority candidates, with contact-pattern-priority candidates highlighted.

Candidate 9 produced the most favourable redocking score at −10.690 kcal mol^-1^, followed by candidates 64 and 83 at −10.060 and −9.860 kcal mol^-1^, respectively. Benchmark redocking gave scores of −8.340 kcal mol^-1^ for RM5 and −9.068 kcal mol^-1^ for baicalin. All 16 report-priority candidates scored more favourably than RM5, and 14 scored more favourably than baicalin under the same docking protocol (Figure 7A, C). However, docking score was not used as the sole selection criterion, because high scores could also reflect increased molecular size or hydrophobicity. The final ranking therefore incorporated QED and other physicochemical properties together with pocket-contact preservation (Figure 7D).

Contact-profile analysis further distinguished high-scoring candidates from candidates that reproduced the RM5-defined interaction environment. Candidates 24, 57, and 29 each recovered 83.3% of the RM5-contact residues, exceeding the recovery observed for the tested natural-product controls. Candidate 24 showed the highest Jaccard similarity to the RM5 contact profile at 0.833. Candidate 9 recovered 72.2% of the reference contacts despite having the strongest docking score. Candidate 57 combined a favourable redocking score with substantial contact-pattern recovery and an interpretable pocket-bound pose; its predicted interaction profile and computationally proposed synthetic route are shown in Figure 8.

**Figure 8.**
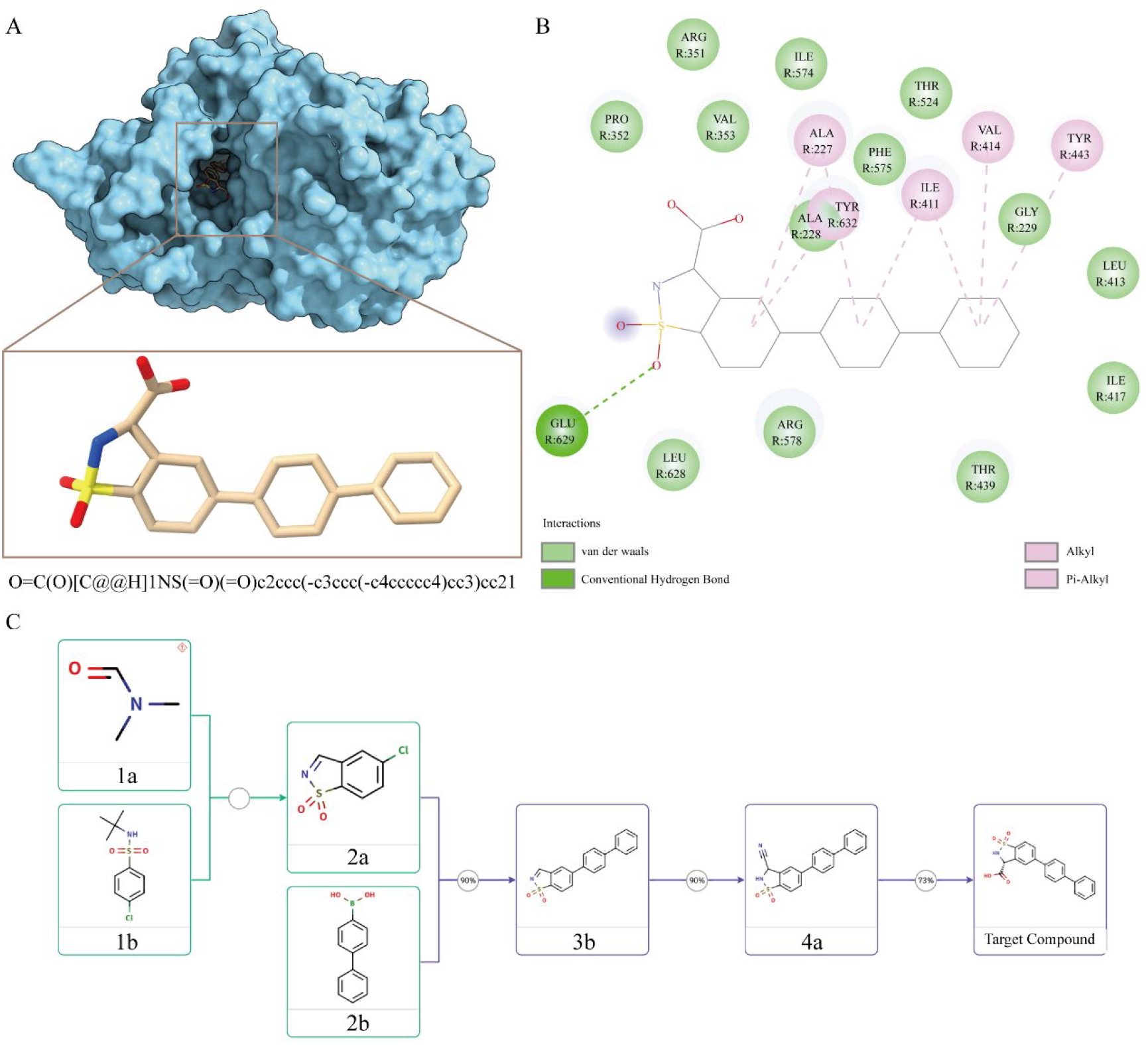
Predicted binding mode, interaction profile, and computationally proposed synthetic analysis of NLRP3_design_57. **Note:** (A) Predicted binding pose of NLRP3_design_57 within the RM5-defined NLRP3 pocket. The protein is shown as a molecular surface and the designed ligand is shown in stick representation. (B) Two-dimensional protein–ligand interaction profile of NLRP3_design_57, showing hydrogen-bond, van der Waals, alkyl, and π-related interactions with surrounding pocket residues. (C) Computationally proposed synthetic route of NLRP3_design_57 based on retrosynthetic analysis, illustrating potential intermediate structures for future synthesis evaluation.

Overall, the RM5-guided workflow reduced 100 generated structures to 16 report-priority NLRP3 candidates. Candidate 9 represented the strongest docking candidate, whereas candidates 24, 57, and 29 provided a better balance between docking performance and preservation of the reference pocket-contact pattern. These candidates formed a focused set for subsequent molecular dynamics simulations, synthesis assessment, and experimental validation.

#### 3.3.2 XM1-pocket-guided generation and prioritisation of AKT1 small-molecule candidates

The XM1-defined binding pocket of AKT1 was used to generate 100 candidate protein–ligand complexes. Sequential ligand-level quality control, PAINS screening, and pocket-geometry inspection retained 88 candidates for redocking. These candidates showed acceptable pocket occupancy, protein–ligand contact geometry, and structural completeness.

All 88 candidates were successfully redocked into the XM1-defined pocket using the same AKT1 receptor preparation, docking box, and AutoDock Vina parameters. Their mode-1 Vina scores ranged from −12.120 to −8.081 kcal mol^-1^. Among them, 31 candidates were classified as very strong, 38 as strong, and 19 as good (Figure 9A, B). Integrated assessment of docking scores, physicochemical properties, structural quality, and pocket-contact features identified 25 report-priority candidates. A further 10 candidates required physicochemical review, 30 were retained for docking-only review, and 23 were not selected.

**Figure 9.**
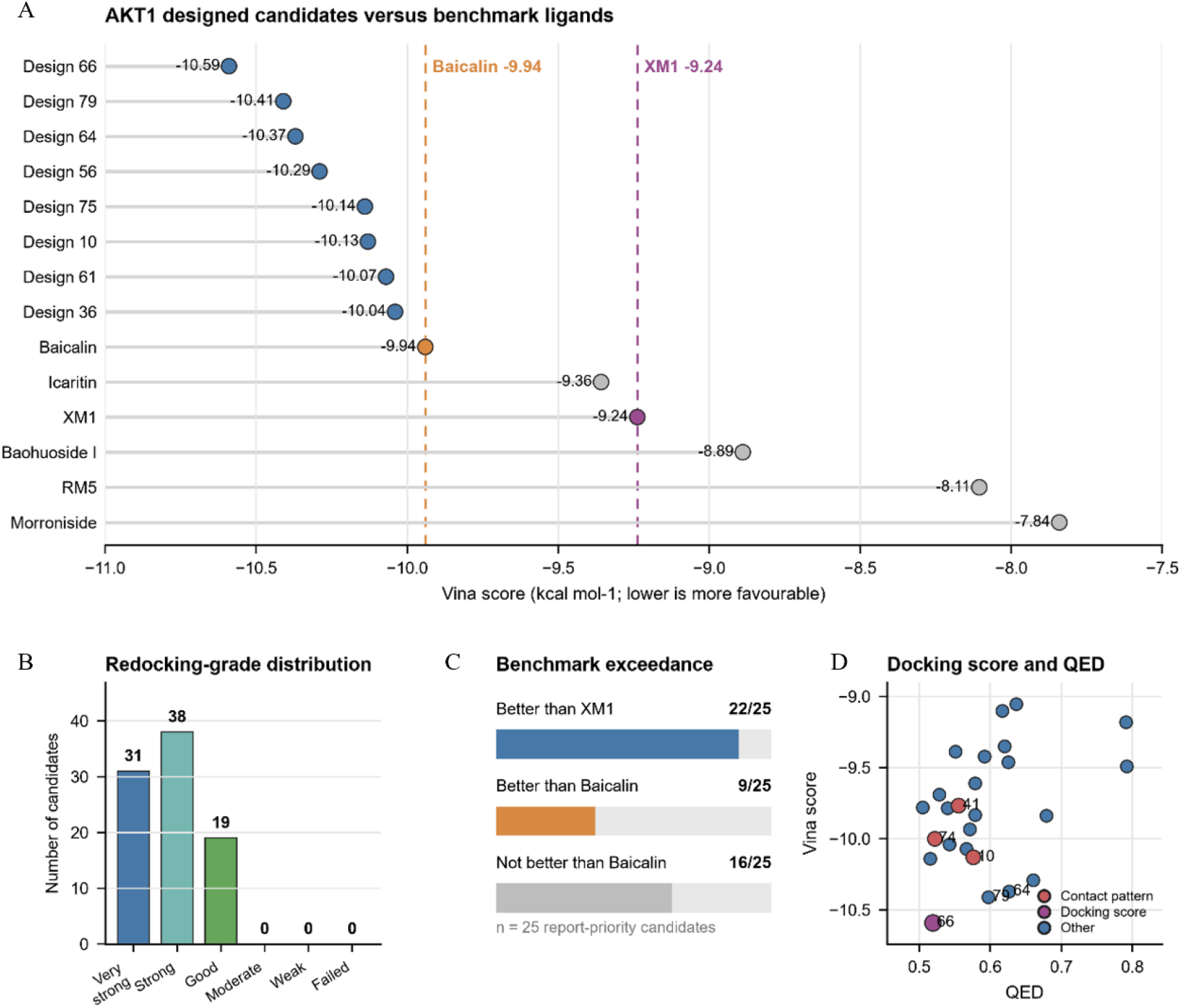
Redocking-based prioritisation of AKT1 small-molecule candidates in the XM1-defined binding pocket. **Note:** (A) Mode-1 Vina scores of representative AKT1-designed candidates and benchmark ligands redocked into the XM1-defined pocket. Dashed lines indicate XM1 and baicalin reference scores. (B) Distribution of redocking-score categories among the final AKT1 candidates. (C) Comparison of report-priority candidates with XM1 and baicalin benchmarks. (D) Relationship between QED values and Vina scores for report-priority candidates, with contact-pattern and docking-score-priority candidates highlighted.

AKT1_design_34 produced the most favourable overall docking score at −12.120 kcal mol^-1^, but it was excluded from the report-priority set because its molecular weight exceeded the predefined range. Among the report-priority candidates, AKT1_design_66 showed the most favourable score at −10.590 kcal mol^-1^, followed by AKT1_design_79, AKT1_design_64, AKT1_design_56, and AKT1_design_10, with scores of −10.410, −10.370, −10.290, and −10.130 kcal mol^-1^, respectively. This result showed that a favourable docking score alone was insufficient for prioritisation when the associated physicochemical properties were less suitable.

Benchmark redocking produced scores of −9.238 kcal mol^-1^ for XM1 and −9.940 kcal mol^-1^ for baicalin. Of the 25 report-priority candidates, 22 scored more favourably than XM1, whereas nine scored more favourably than baicalin under the same docking protocol (Figure 9A, C). The relationship between QED and docking score further distinguished candidates with strong docking performance from those offering a more balanced physicochemical profile (Figure 9D).

Contact-profile analysis was then used to assess preservation of the XM1-defined binding environment. AKT1_design_41 recovered 19 of the 22 XM1-contact residues, corresponding to 86.4% recovery and a Jaccard similarity of 0.792. AKT1_design_10 and AKT1_design_74 each recovered 81.8% of the reference contacts. By comparison, icaritin and baicalin recovered 81.8% and 59.1%, respectively. AKT1_design_66 and AKT1_design_79 therefore represented docking-score-prioritised candidates, whereas AKT1_design_41, AKT1_design_10, and AKT1_design_74 showed stronger preservation of the XM1-defined contact pattern. The predicted binding mode, interaction profile, and computationally proposed synthetic route of AKT1_design_41 are presented in Figure 10.

**Figure 10.**
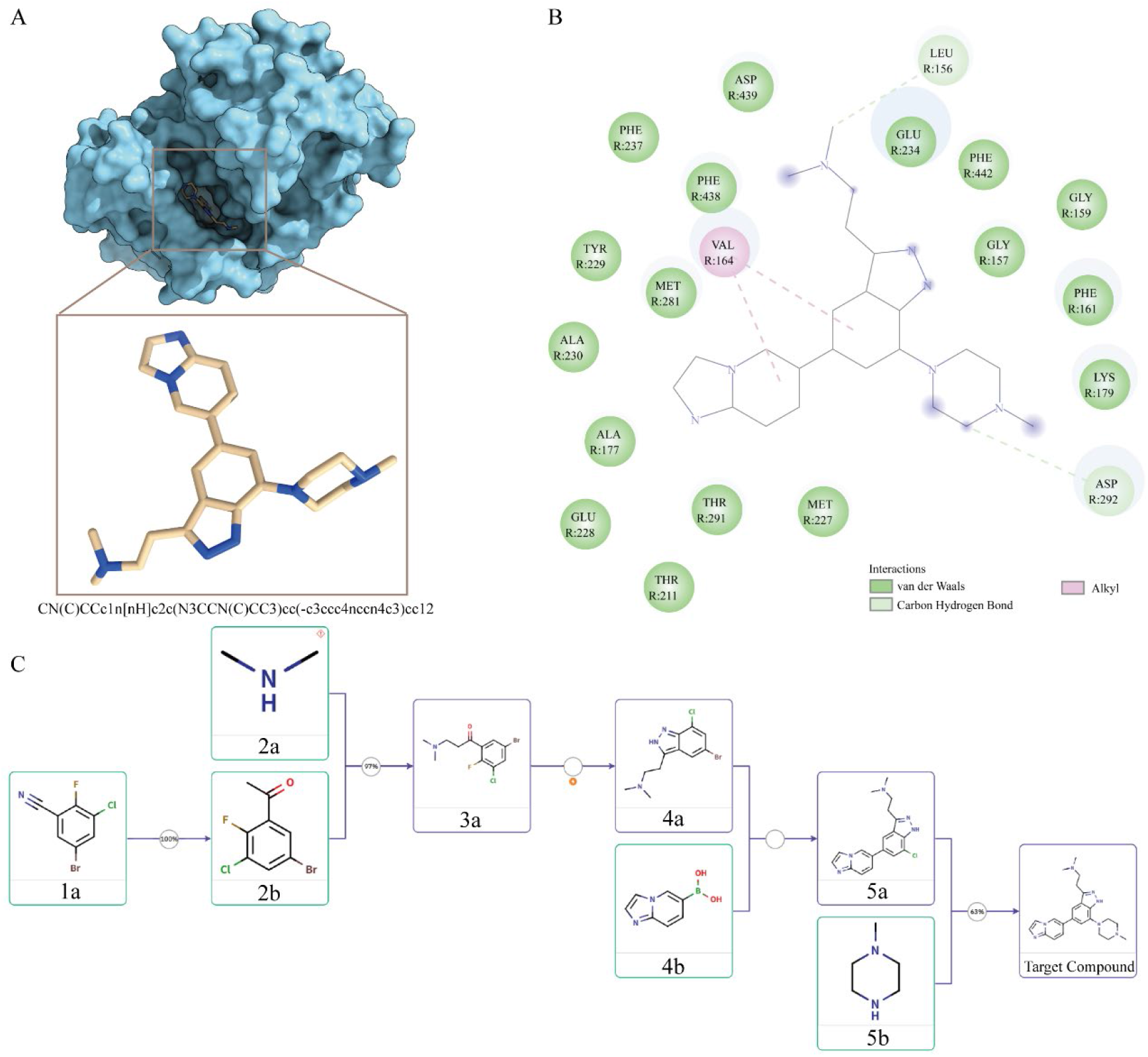
Predicted binding mode, interaction profile, and computationally proposed synthetic analysis of AKT1_design_41. **Note:** (A) Predicted binding pose of AKT1_design_41 within the XM1-defined AKT1 binding pocket. AKT1 is shown as a molecular surface, and the designed ligand is shown in stick representation. The inset shows the enlarged ligand conformation within the predicted binding region. (B) Two-dimensional protein–ligand interaction profile of AKT1_design_41, showing van der Waals, carbon–hydrogen bond, and alkyl interactions with surrounding pocket residues. (C) Computationally proposed synthetic analysis of AKT1_design_41 based on retrosynthetic evaluation, illustrating potential intermediate structures for future synthesis assessment.

Overall, the XM1-guided workflow reduced 100 generated structures to 25 report-priority AKT1 candidates. The selected set included both high-scoring candidates and candidates that more closely reproduced the reference pocket-contact pattern, providing a focused basis for subsequent molecular dynamics simulations, synthesis assessment, and experimental validation.

#### 3.3.3 DF6-pocket-guided generation and prioritisation of MET small-molecule candidates

The DF6-defined kinase pocket of MET was used to generate 100 candidate protein–ligand complexes. All candidates passed the initial ligand-level and complex-level quality-control assessment, and none triggered PAINS alerts. Subsequent inspection of pocket occupancy, protein–ligand contacts, and steric compatibility retained 97 candidates for redocking.

The 97 candidates were successfully redocked into the DF6-defined pocket using the same MET receptor preparation, docking box, and AutoDock Vina parameters. Their mode-1 Vina scores ranged from −12.770 to −7.820 kcal mol^-1^. Of these candidates, 51 were classified as very strong, 29 as strong, 14 as good, and three as moderate (Figure 11A,B). Integrated assessment of docking performance, physicochemical properties, structural quality, and pocket-contact features identified 43 report-priority candidates. A further 16 candidates required physicochemical review, 21 were retained for docking-only review, and 17 were not selected.

**Figure 11.**
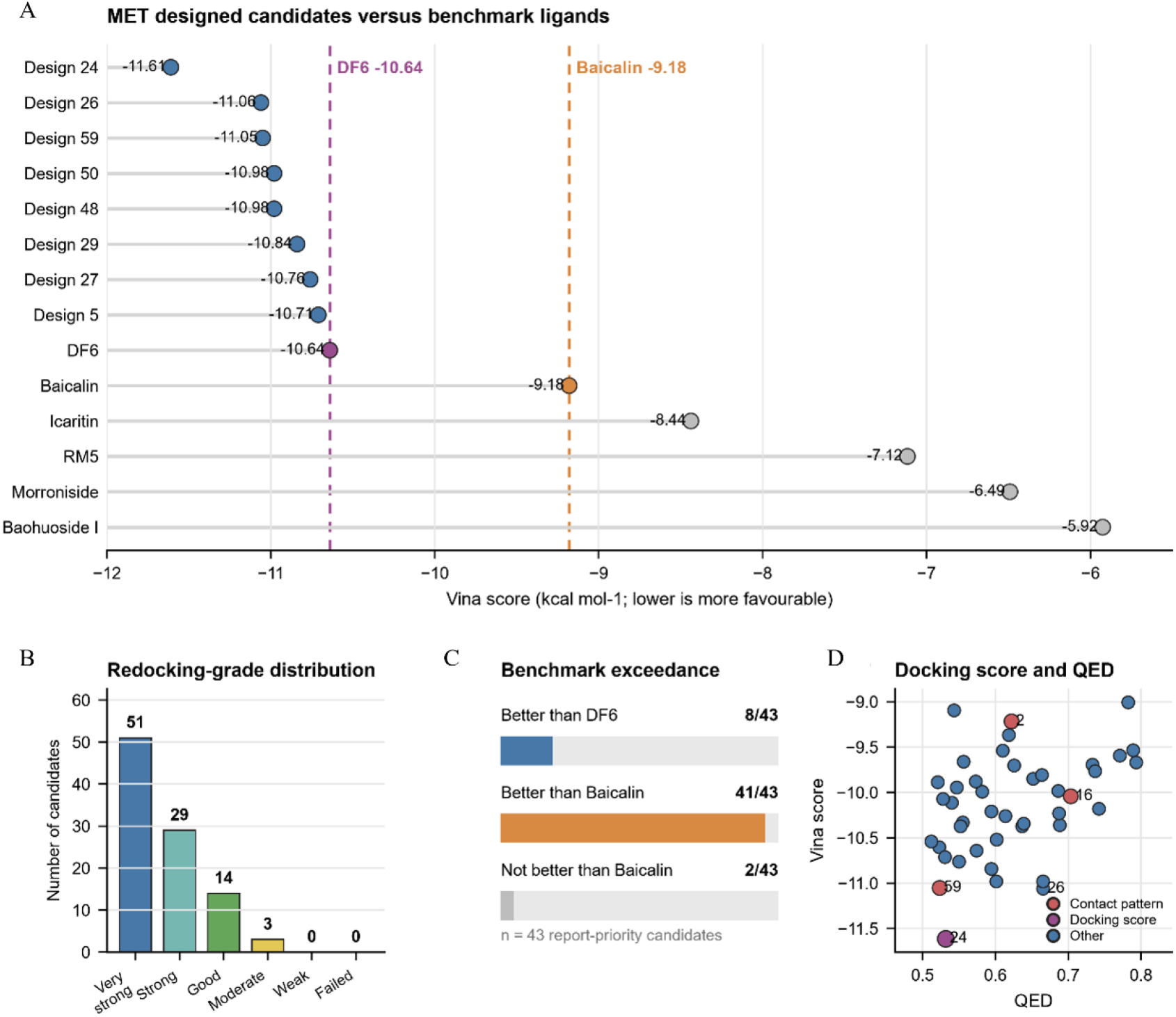
Redocking-based prioritisation of MET small-molecule candidates in the DF6-defined binding pocket. **Note:** (A) Mode-1 Vina scores of representative MET-designed candidates and benchmark ligands redocked into the DF6-defined pocket. Dashed lines indicate DF6 and baicalin reference scores. (B) Distribution of redocking-score categories among the final MET candidates. (C) Comparison of report-priority candidates with DF6 and baicalin benchmarks. (D) Relationship between QED values and Vina scores for report-priority candidates, with contact-pattern and docking-score-priority candidates highlighted.

MET_design_31 produced the most favourable overall docking score at −12.770 kcal mol^-1^, followed by MET_design_54 and MET_design_38 at −12.440 and −12.430 kcal mol^-1^, respectively. However, these candidates were excluded from the report-priority set because of low QED values, elevated LogP, or both. Among the report-priority candidates, MET_design_24 showed the strongest balanced profile, with a docking score of −11.610 kcal mol^-1^, a QED value of 0.531, and acceptable pocket-contact characteristics. Other highly ranked candidates included MET_design_26, MET_design_59, MET_design_50, and MET_design_48, with scores ranging from −11.060 to −10.980 kcal mol^-1^. These results further showed that docking score alone was insufficient for final candidate selection.

Benchmark redocking produced a score of −10.640 kcal mol^-1^ for DF6 and −9.180 kcal mol^-1^ for baicalin. Among the 43 report-priority candidates, eight scored more favourably than DF6 and 41 scored more favourably than baicalin under the same docking protocol (Figure 11 A, C). The relationship between QED and docking score further separated candidates with strong docking performance from those with more balanced physicochemical properties (Figure 11 D).

Contact-profile analysis was used to assess preservation of the DF6-defined binding environment. MET_design_59, MET_design_2, and MET_design_16 each recovered 91.7% of the DF6-contact residues. MET_design_2 showed the highest Jaccard similarity to the DF6 contact profile at 0.917, whereas MET_design_59 combined high contact recovery with report-priority docking performance. MET_design_24 recovered 87.5% of the reference contacts and showed a Jaccard similarity of 0.875. Its predicted binding mode, interaction profile, and computationally proposed synthetic route are presented in Figure 12.

**Figure 12.**
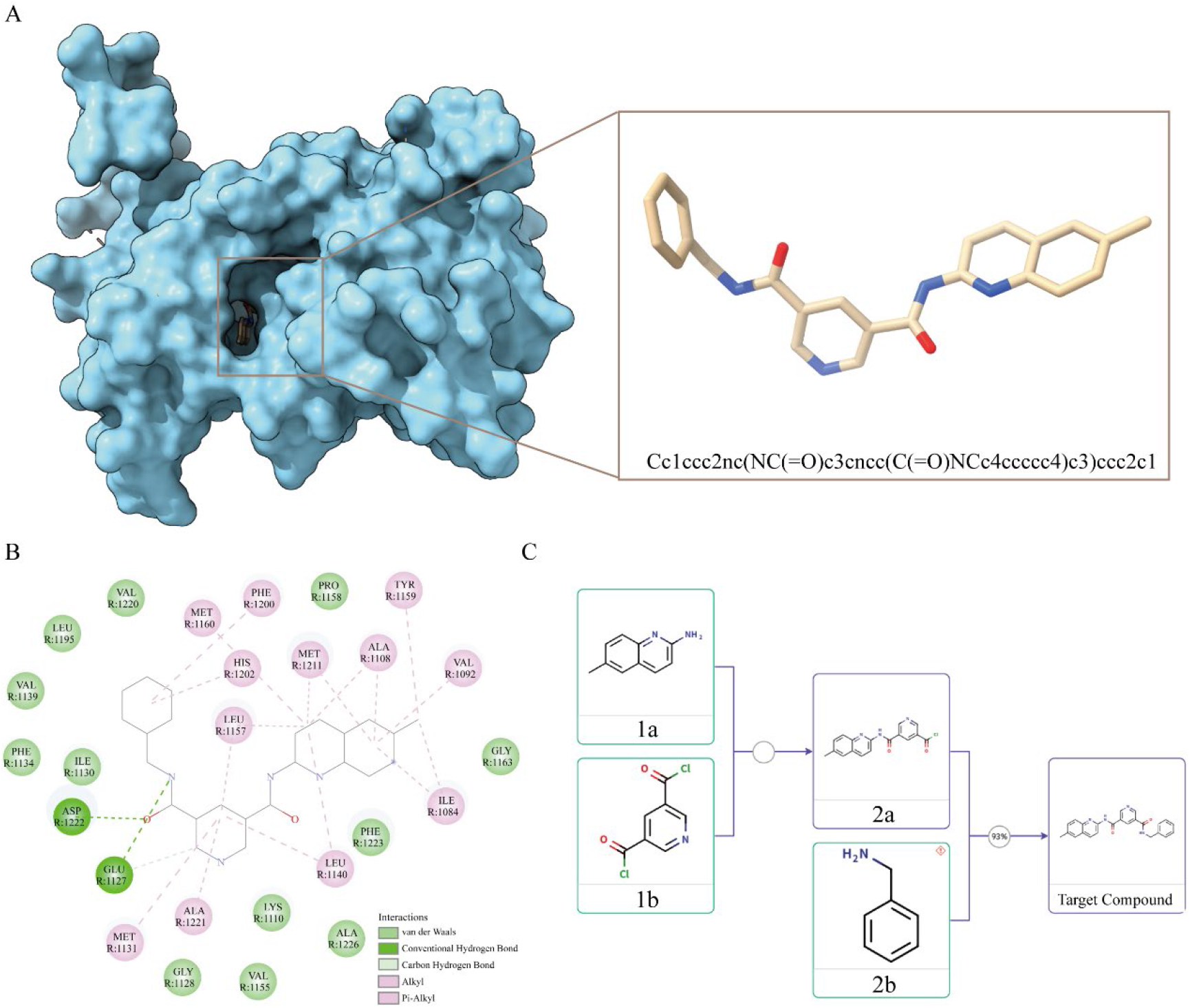
Predicted binding mode, interaction profile, and computationally proposed synthetic analysis of MET_design_24. **Note:** (A) Predicted binding pose of MET_design_24 within the DF6-defined MET binding pocket. MET is shown as a molecular surface and the designed ligand is shown in stick representation. The inset shows the enlarged ligand conformation within the predicted binding region. (B) Two-dimensional protein–ligand interaction profile of MET_design_24, showing van der Waals, hydrogen-bond-related, carbon–hydrogen bond, alkyl, and π-related interactions with surrounding pocket residues. (C) Computationally proposed synthetic analysis of MET_design_24 based on retrosynthetic evaluation, illustrating potential intermediate structures for future synthesis assessment.

Overall, the DF6-guided workflow reduced 100 generated structures to 43 report-priority MET candidates. The selected set included both docking-score-prioritised molecules and candidates that more closely reproduced the reference ligand-contact pattern. These candidates provide a focused basis for subsequent molecular dynamics simulations, synthesis assessment, and experimental validation. This kinase-pocket design strategy is distinct from the antibody-like binder workflow targeting the extracellular region of MET for potential EV recognition.

#### 3.3.4 ID8-pocket-guided generation and prioritisation of PTGS2 small-molecule candidates

The ID8-defined binding pocket of PTGS2 was used to generate 50 candidate protein–ligand complexes. Initial ligand-level and complex-level quality control identified 47 candidates as passing and three as requiring review, with no PAINS alerts detected. Subsequent inspection of pocket occupancy, protein–ligand contacts, and steric compatibility retained 43 candidates for redocking.

All 43 candidates were redocked into the ID8-defined pocket using the same PTGS2 receptor preparation, docking box, and AutoDock Vina parameters. Their mode-1 Vina scores ranged from −9.294 to 4.586 kcal mol^-1^. The final set included two strong, 12 good, nine moderate, and 20 weak candidates, with no candidate classified as very strong (Figure 13A, B). This distribution indicated a more selective docking outcome than those observed for the AKT1 and MET design workflows.

**Figure 13.**
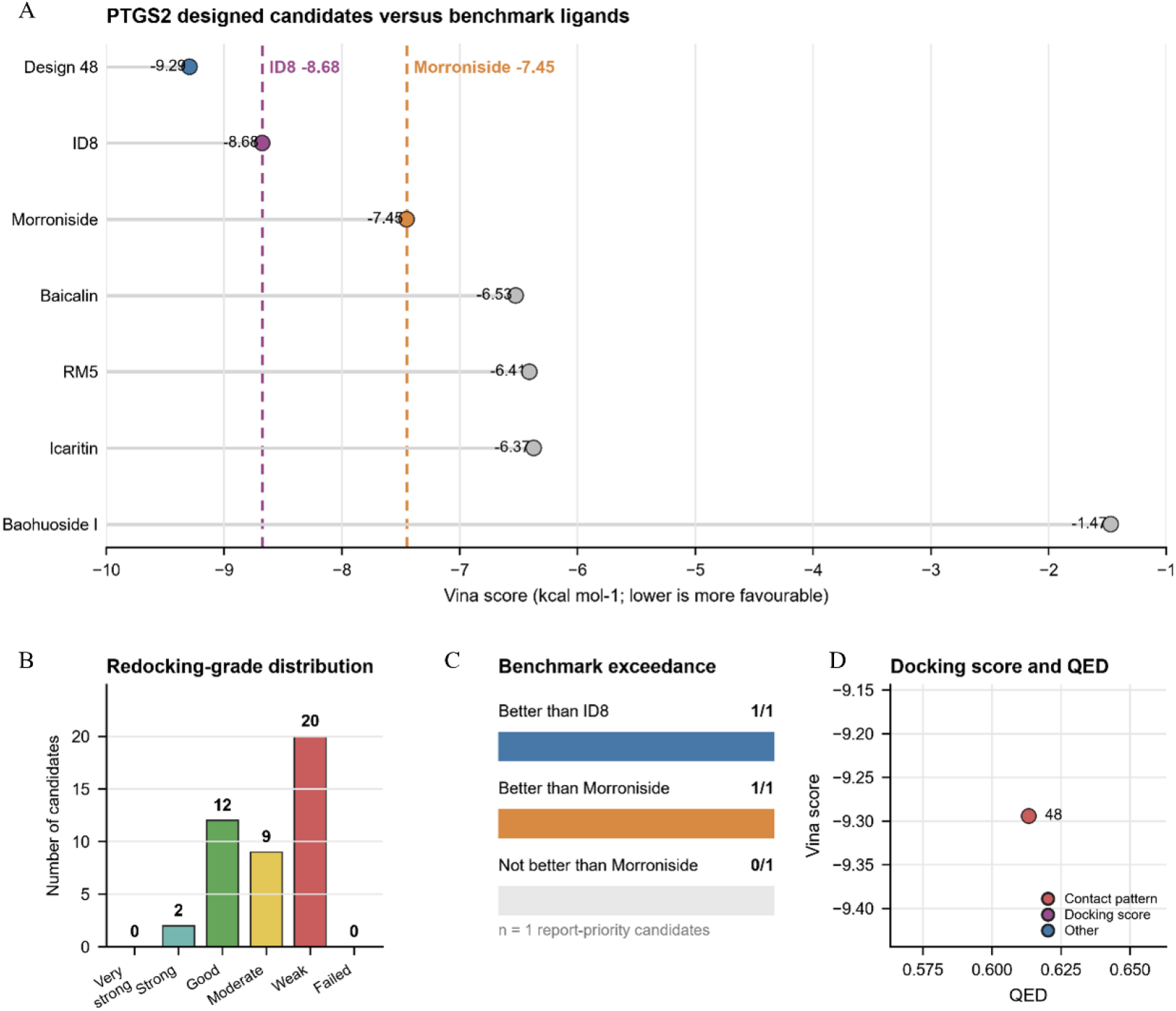
Redocking-based prioritisation of PTGS2 small-molecule candidates in the ID8-defined binding pocket. **Note:** (A) Mode-1 Vina scores of the selected PTGS2-designed candidate and benchmark ligands redocked into the ID8-defined pocket. Dashed lines indicate ID8 and morroniside reference scores. (B) Distribution of redocking-score categories among the final PTGS2 candidates. (C) Comparison of the report-priority candidate with ID8 and morroniside benchmarks. (D) Relationship between QED and Vina score for the report-priority candidate, highlighting the contact-pattern-priority candidate.

Integrated assessment of docking performance, physicochemical properties, structural quality, and pocket-contact features retained only PTGS2_design_48 as a report-priority candidate. It showed a mode-1 Vina score of −9.294 kcal mol^-1^, a QED value of 0.613, a molecular weight of 406.53 Da, a LogP value of 4.799, and a tPSA of 93.89 Å^2^. The candidate contacted 15 pocket residues and showed a ligand contact-atom fraction of 0.867. Other generated molecules were not prioritised because of weaker docking scores, limited pocket-contact characteristics, or less favourable integrated physicochemical profiles.

Benchmark redocking produced a score of −8.677 kcal mol^-1^ for ID8. Among the natural-product controls, morroniside showed the most favourable score at −7.451 kcal mol^-1^, followed by baicalin, RM5, icaritin, and baohuoside I. PTGS2_design_48 therefore scored more favourably than both ID8 and all tested natural-product controls under the same docking protocol (Figure 13A, C).

Contact-profile analysis showed that PTGS2_design_48 recovered 12 of the 14 ID8-contact residues, corresponding to 85.7% recovery and a Jaccard similarity of 0.600. Morroniside recovered the complete ID8 contact set but showed a weaker docking score and a less favourable physicochemical profile. Baohuoside I also recovered 85.7% of the reference contacts but showed substantially weaker redocking performance. PTGS2_design_48 was therefore prioritised because it provided the best overall balance among docking performance, physicochemical properties, and preservation of the ID8-defined interaction environment. Its predicted binding mode, interaction profile, and computationally proposed synthetic route are presented in Figure 14.

**Figure 14.**
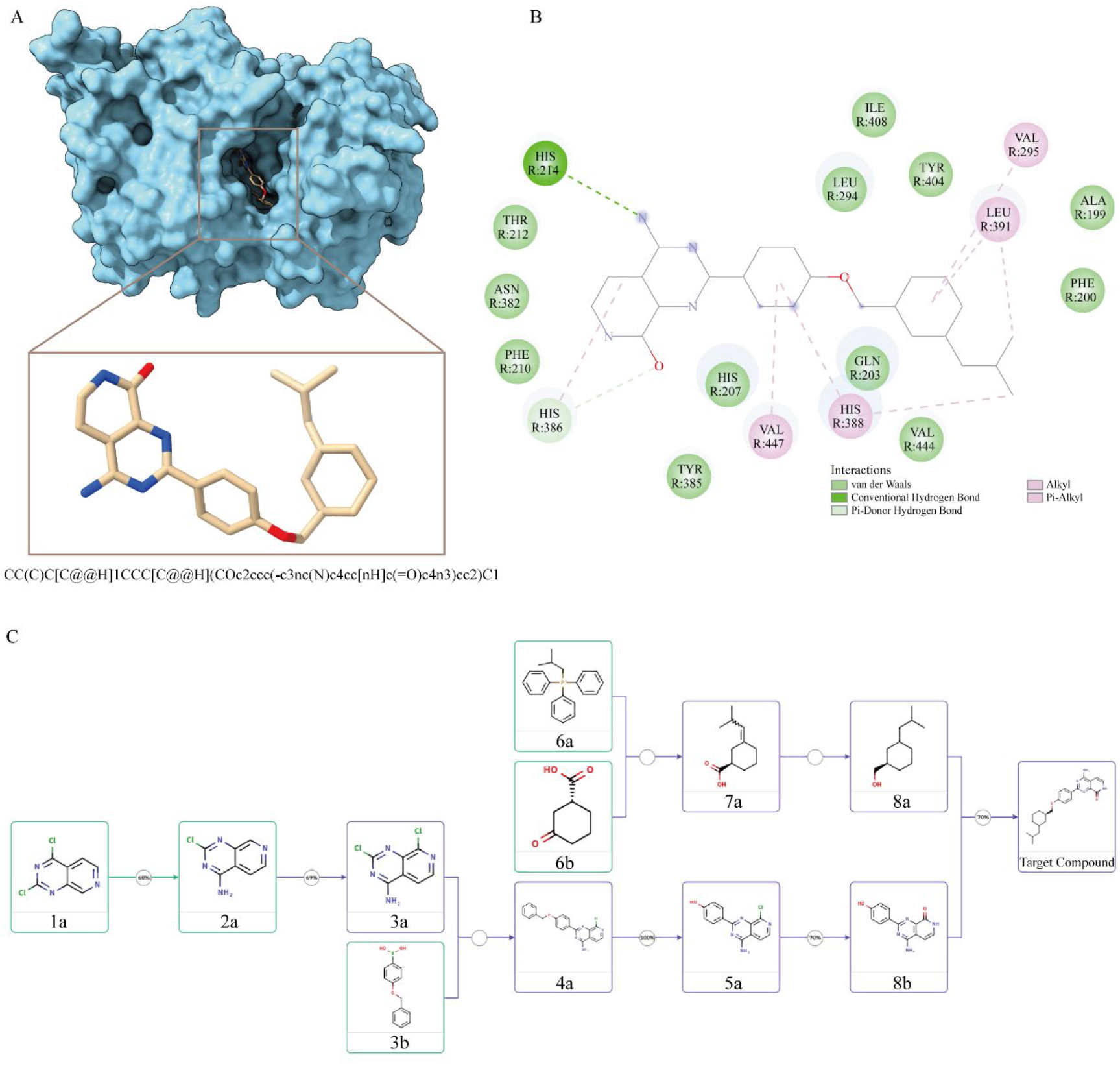
Predicted binding mode, interaction profile, and computationally proposed synthetic analysis of PTGS2_design_48. **Note:** (A) Predicted binding pose of PTGS2_design_48 within the ID8-defined PTGS2 binding pocket. PTGS2 is shown as a molecular surface and the designed ligand is shown in stick representation. The inset shows the enlarged ligand conformation within the predicted binding region. (B) Two-dimensional protein–ligand interaction profile of PTGS2_design_48, showing hydrogen-bond-related, van der Waals, alkyl, and π-related interactions with surrounding pocket residues. (C) Computationally proposed synthetic analysis of PTGS2_design_48 based on retrosynthetic evaluation, illustrating potential intermediate structures for future synthesis assessment.

Overall, the ID8-guided workflow reduced 50 generated structures to a single report-priority candidate. PTGS2_design_48 represents a selectively prioritised molecule for subsequent molecular dynamics simulations, synthesis assessment, and experimental validation.

### 3.4 Computational design and prioritisation of antibody-like binders targeting CD81 and MET

VHH and scFv candidates targeting CD81 and MET were computationally designed and evaluated using a unified structural prioritisation framework to identify potential EV-recognition or EV-capture binders. For CD81, two reference complexes, PDB 5DFW and PDB 6U9S, were used to generate antibody-like binder candidates. A total of 40 CD81-directed designs were produced, of which eight candidates were retained after integrated structural and developability-based filtering. The remaining candidates were excluded mainly due to incomplete model-confidence information, insufficient hotspot recovery, unfavourable interface geometry, or severe atomic clashes.

The 5DFW-derived CD81 design set produced CD81-scFv n=3 as a balanced candidate across interface recovery, structural confidence, and sequence-level properties. This candidate recovered all six predefined hotspot residues and formed contacts with 26 antigen residues within 4.5 Å, while exhibiting two severe clashes. Among candidates with available co-folding metrics, CD81-scFv n=3 showed the strongest interface-confidence profile within this design group, with an iPTM value of 0.518 and an ipAE value of 15.15. Its PRODIGY-derived interaction estimate corresponded to a predicted binding free energy of −10.2 kcal mol^-1^, with a predicted K_d_ estimate of 3.2 × 10^-8^ M. Sequence-based developability assessment predicted a heavy-chain solubility of 57.7%, together with favourable predicted degradation-related parameters. Based on the combined evaluation of hotspot recovery, interface characteristics, structural confidence, and developability-related features, CD81-scFv n=3 was selected as a structure-balanced representative candidate.

The 6U9S-derived CD81 design set produced a different candidate profile. CD81-VHH n=2 showed stronger predicted interaction characteristics but higher structural risk. This candidate recovered all six hotspot residues and formed contacts with 31 antigen-interface residues. Its predicted binding free energy was −12.5 kcal mol^-1^, corresponding to a predicted K_d_ estimate of 6.6 × 10^-1^⁰ M. However, the model showed limited co-folding confidence, with an iPTM value of 0.150 and an ipAE value of 23.95, together with nine severe clashes approaching the predefined exclusion threshold. Therefore, CD81-VHH n=2 was classified as an affinity-prioritised computational candidate requiring further interface optimisation, whereas CD81-VHH n=4 represented a lower-clash alternative candidate.

The MET-directed design sets showed a distinct candidate profile from CD81. MET-VHH and MET-scFv candidates were generated using the same MET reference structure but different binder architectures. Within the MET-VHH design set, 17 of 20 candidates passed geometric filtering, whereas three were excluded because they failed to recover sufficient hotspot residues. MET-VHH n=14 (Figure 15) was prioritised based on integrated interface evaluation. This candidate recovered four of six hotspot residues, formed contacts with 30 antigen-interface residues, and contained one severe clash. Its predicted binding free energy was −13.5 kcal mol^-1^, corresponding to a predicted K_d_ estimate of 1.3 × 10^-1^⁰ M. Sequence-level evaluation predicted a heavy-chain solubility of 63.8%. These results indicate that MET-VHH n=14 represents a structurally favourable computational candidate for subsequent experimental testing.

**Figure 15.**
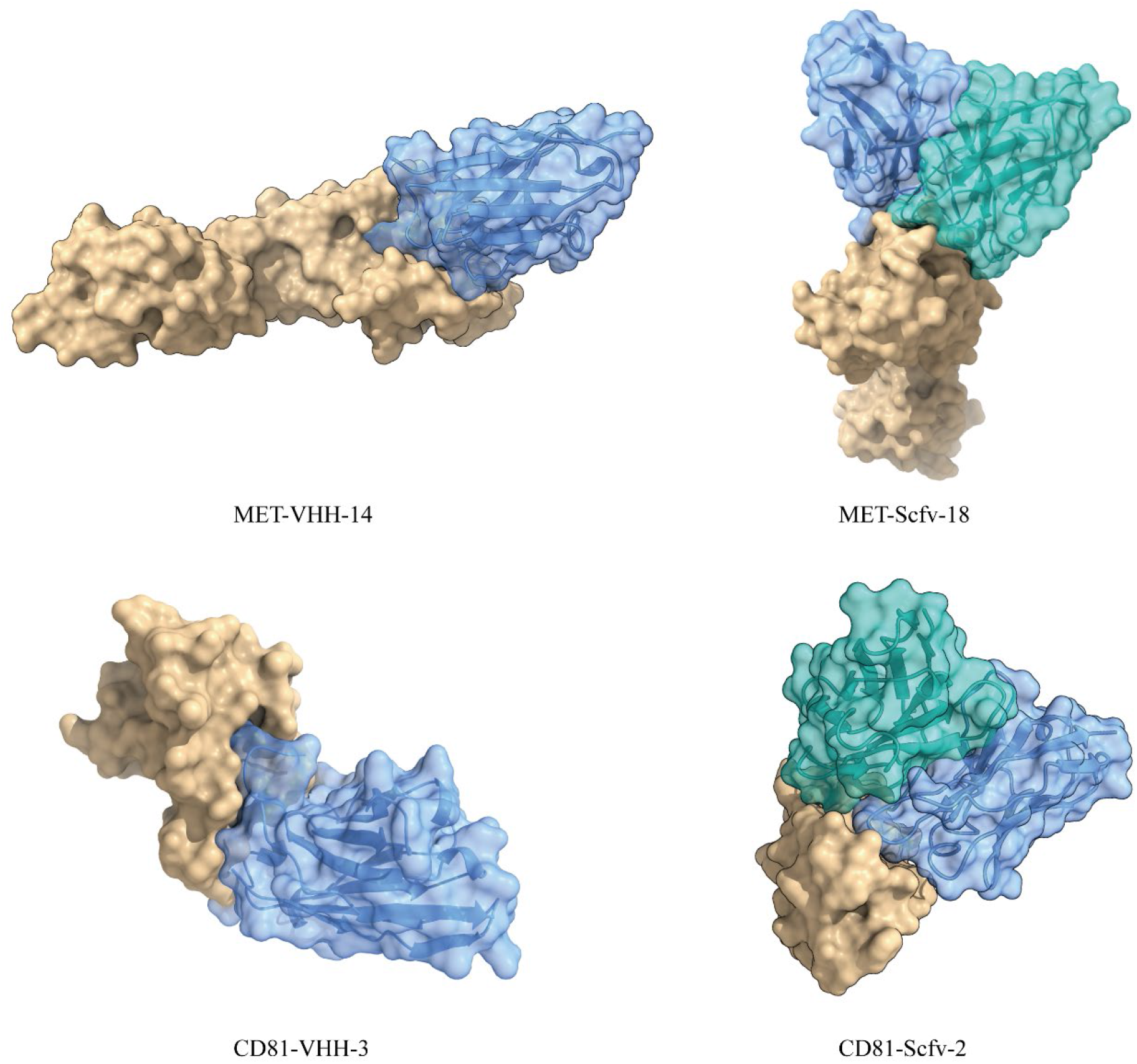
Predicted binding modes of computationally designed VHH and scFv binders targeting MET and CD81. **Note:** Representative predicted complexes of antibody-like binders targeting MET and CD81. Antigen molecules are shown as light-brown molecular surfaces, whereas designed binders are shown as semi-transparent blue or cyan molecular surfaces with cartoon representations. MET-VHH-14 and CD81-VHH-3 represent VHH-like single-domain binders, whereas MET-scFv-18 and CD81-scFv-2 represent scFv-like binders containing paired variable domains. The predicted models illustrate candidate binder orientations and antigen-contact interfaces for potential MET- or CD81-associated EV recognition applications.

Within the MET-scFv design set, 12 of 20 candidates were retained after structural filtering. Several designs with complete co-folding metrics were excluded because of low interface confidence, elevated ipAE values, or excessive clashes. Among retained candidates, MET-scFv n=18 showed the most favourable interface geometry. This candidate recovered all six hotspot residues, formed contacts with 19 antigen-interface residues, and showed no severe clashes. Its predicted binding free energy was −12.3 kcal mol^-1^, corresponding to a predicted K_d_ estimate of 9.0 × 10^-1^⁰ M. Predicted solubility values for the heavy and light chains were 55.4% and 61.9%, respectively. MET-scFv n=18 was therefore prioritised as a low-clash candidate with complete hotspot recovery, although further experimental evaluation is required.

Taken together, the CD81 and MET design results revealed different computational candidate profiles. CD81-scFv n=3 represented a structure-balanced candidate with stronger model-confidence support, whereas CD81-VHH n=2 represented a higher predicted-interaction but higher-risk design. MET-VHH n=14 and MET-scFv n=18 represented the leading candidates from their respective design strategies, combining favourable interface characteristics with acceptable structural features. These antibody-like binders provide computationally prioritised candidates for subsequent expression, binding assays, and EV-recognition evaluation.

## 4 Discussion

PDAC is one of the most aggressive malignancies, and current treatments rarely produce durable responses. This difficulty does not arise solely from the tumour cells themselves. It is also shaped by the complex communication among tumour cells, stromal components, and immune-cell populations within the tumour microenvironment [37]. Against this background, the present study developed a computational framework for prioritising candidates that may interfere with PDAC EV-associated immune regulation from two complementary directions. The production-side strategy aims to modulate EV-producing cells and reduce the generation or release of harmful vesicles, whereas the action-side strategy seeks to recognise or intercept EVs after they have already entered the extracellular environment.

The study first established a literature-curated framework containing 88 genes associated with PDAC EV biology. These genes covered EV biogenesis and release, cargo loading, immune escape, and remodelling of the tumour microenvironment. They were then compared with the cell-type-resolved prognostic annotations available in the ctPANDA database [25], yielding 42 candidate genes supported by both EV-related mechanistic evidence and clinical prognostic associations. On this basis, a library of 18 natural products derived from *Scutellaria baicalensis*, *Epimedium* spp., and *Cornus officinalis* was assembled, and their potential human protein targets were predicted using SwissTargetPrediction. Intersecting these predicted targets with the 42 ctPANDA-supported genes identified 13 PDAC EV-related targets that might be covered by the natural-product library. Seven representative compounds were then selected for molecular docking according to target coverage, balanced representation of the three plant sources, and chemical diversity. Five protein targets were selected from the 13 overlapping candidates because they were relevant to PDAC EV-associated immune regulation, were predicted to be affected by the natural products, and had experimentally resolved ligand-bound structures. Through this stepwise prioritisation, a broad initial candidate space was narrowed to a focused group of compounds and targets supported by mechanistic relevance, clinical association, predicted chemical coverage, and structural tractability.

Based on these prioritised targets, the study further explored natural compounds that might regulate the biogenesis of tumour-derived EVs and thereby limit tumour progression. Previous studies suggest that certain natural products can interfere with EV-related biological processes [38–40]. However, natural products often have practical limitations, including modest potency, limited selectivity, poor physicochemical properties, and restricted opportunities for structural optimisation. To address these issues, experimentally resolved ligand-binding pockets were used to guide de novo small-molecule generation, followed by structural and physicochemical screening. This approach was intended to complement the natural-product candidates. Nevertheless, computational prioritisation remains only a starting point. Whether the selected natural products can reduce EV release or alter immunosuppressive cargo without causing substantial tumour-cell damage must still be tested experimentally. At least two PDAC cell lines with distinct molecular backgrounds should therefore be examined, together with a non-malignant pancreatic ductal epithelial-cell line as a control.

Baicalin and icaritin may serve as the first natural-product candidates for experimental validation. Both showed favourable docking results against targets including PTGS2, AKT1, MET, and NLRP3, but docking alone indicates only potential compatibility and cannot establish direct binding or functional regulation. Concentration-response and time-course experiments should therefore be used to define a non-cytotoxic treatment range. Metabolic activity, membrane integrity, apoptosis, proliferation, and viable-cell number should be assessed in parallel [41]. This step is essential because a reduction in EV abundance may simply reflect a loss of viable producer cells rather than direct inhibition of EV biogenesis or secretion. EV output should therefore be normalised to viable-cell number and collection time so that general cytotoxicity is not mistaken for specific regulation of EV production.

Subsequent validation should reflect the biological function of each proposed target. PTGS2-related activity may be assessed through COX-2 enzyme-activity assays or measurements of prostaglandin production, with an established COX-2 inhibitor included as a benchmark [42]. AKT1- and MET-related effects may be evaluated by measuring phosphorylation and downstream signalling changes. NLRP3-related activity may be examined through complementary inflammasome readouts, including ASC-speck formation, caspase-1 cleavage, mature IL-1β release, and gasdermin D cleavage. Where feasible, biolayer interferometry, microscale thermophoresis, or thermal-shift analysis may be used to determine whether a compound directly binds the proposed target protein. The strongest compound–target pairs should then be examined using target knockdown, knockout, or expression-rescue experiments [43]. If the EV-related effect weakens or is restored when the target is altered, the evidence for a target-dependent mechanism becomes considerably stronger. Only candidates that show reproducible target-related activity within a non-cytotoxic range, and whose effects on EVs remain evident after viable-cell normalisation, should proceed to immune-functional and combination studies.

Whereas the production-side strategy aims to reduce the continuous release of harmful EVs, the action-side strategy addresses a different but equally important question: whether EVs that have already entered the extracellular space can be recognised and intercepted. MET and CD81 were selected because they serve distinct but complementary roles within this framework. Previous studies have reported MET as a tumour-associated surface component on EVs in pancreatic cancer-related settings, suggesting that it may help identify, capture, or block specific tumour-associated EV subpopulations [44, 45]. However, MET expression alone cannot confirm that a detected particle is an EV. Biological samples may also contain soluble MET, MET-bearing membrane fragments, protein aggregates, or other non-vesicular particles. CD81, by contrast, is a widely used EV-associated tetraspanin marker [46]. A sequential recognition strategy may therefore be more reliable. MET could first be used to enrich particles that are more likely to originate from tumour cells, after which CD81 could be detected to determine whether those particles also display a typical EV-associated surface feature. This approach may reduce false-positive signals arising from soluble MET, membrane debris, or other non-EV material, while providing a possible route for intercepting EV-mediated communication that supports tumour growth, metastasis, and immune escape. VHHs and scFvs offer different advantages for this purpose. VHHs are compact and generally stable and can often be produced efficiently in *Escherichia coli* [47]. scFvs contain paired VH and VL domains and may also be expressed in mammalian systems as humanised Fc-fusion constructs, which can improve stability, circulating half-life, and potential effector functions [14].

Before intact EVs are examined, the reliability of the VHH and scFv candidates themselves must be established. Each candidate should be expressed and purified reproducibly and should remain predominantly monomeric. Size-exclusion chromatography can be used to assess monomer content and aggregation [48], whereas nano-differential scanning fluorimetry or differential scanning calorimetry can be used to evaluate thermal stability [49]. Surface plasmon resonance or biolayer interferometry may then be used to measure the affinity and binding kinetics of each candidate against the MET extracellular domain or the large extracellular loop of CD81 [50]. These experiments should include appropriate concentration series, irrelevant proteins, irrelevant VHH or scFv controls, and a reference channel. Once binding has been confirmed, site-directed mutagenesis or alanine scanning may be used to identify residues that make important contributions to the predicted epitope [51]. Target specificity is equally important. MET-directed candidates should be compared across MET-high, MET-low, and MET-knockout cells. Ideally, binding should decrease as MET expression falls and should become minimal after MET knockout. CD81-directed candidates should be assessed in the same way, while also being tested for cross-reactivity against CD9, CD63, and unrelated membrane proteins.

Only after protein- and cell-level validation should the candidates be tested against intact EVs. EV isolation and characterisation should follow the main recommendations of MISEV2023 [52, 53], using complementary measurements such as nanoparticle tracking analysis, transmission or cryogenic electron microscopy, and EV-associated protein markers. EVs may be collected from MET-high and MET-low PDAC cell lines, with non-malignant pancreatic ductal epithelial cells included as a control. MET- or CD81-knockout cells would provide especially informative controls where available. EV inputs should be normalised by particle number. CD81-directed candidates may initially be evaluated as EV-capture reagents using bead-based flow cytometry, plate-based immunoassays, or single-particle platforms. Captured material should then be tested for additional EV-associated markers such as CD9 and CD63 to confirm that it has an EV-like phenotype. MET-directed candidates should preferentially recognise EVs derived from MET-high PDAC cells, with weaker binding to EVs from MET-low, MET-deficient, or non-malignant cells. Competition with soluble MET may provide further evidence of target-dependent binding. If EV interception or uptake inhibition is examined, dye-only controls, irrelevant binder controls, and an independent labelling method should also be included. A reduction in fluorescence does not necessarily indicate reduced EV uptake because aggregation, lower EV recovery, or altered dye behaviour can produce a similar result. Candidate selection should therefore not be based on affinity alone. Expression yield, purity, monomer content, specificity, recognition of intact EVs, aggregation tendency, and storage stability all need to be considered together. The most promising candidate is likely to be the one that achieves the best overall balance rather than the one with the strongest single measurement.

Once the production-side small molecules and post-release VHH or scFv candidates have each shown reliable activity, their combined effects can be examined to determine whether the two intervention layers may produce an effect greater than either alone. A 2 × 2 factorial designs would provide a suitable starting point. PDAC cells would first receive vehicle or a candidate small molecule, after which released EVs would be collected and incubated with a target-specific VHH, scFv, irrelevant binder, or buffer before being added to recipient immune cells. The central groups would include control EVs, EVs produced after small-molecule treatment, control EVs treated with a VHH or scFv, and EVs exposed to both interventions. EV-depleted conditioned medium, binder-only controls, compound-carryover controls, and equal-particle-input controls should also be included. These controls would help separate the effects of soluble factors, the binder itself, and residual small molecules. Equal-particle-input comparisons are particularly important because they distinguish a reduction in total EV output from a change in the immunosuppressive activity of each individual EV. Primary human monocyte-derived macrophages may be used as the main recipient-cell model, with THP-1-derived macrophages included as a more standardised complementary system [54]. Relevant readouts include EV uptake, cell viability, cytokine secretion, phagocytic activity, and the expression of CD163, CD206, HLA-DR, and CD86 [55]. Macrophage responses should not be reduced to a simple M1/M2 classification. Surface phenotype, secretory behaviour, and functional activity should instead be interpreted together. Activated primary human CD8⁺ T cells may be used to assess adaptive immune effects, including proliferation, viability, IFN-γ production, granzyme B expression, degranulation, and tumour-cell killing. PD-1 may be recorded as a supporting phenotypic marker, but it cannot by itself establish T-cell exhaustion. If PD-L1-positive EVs appear to contribute to immune suppression, mechanistic controls should include PD-L1-blocking antibodies, PD-L1-deficient EVs, or EV preparations depleted of PD-L1-positive particles.

An enhanced response in the combined group would initially support an additive or complementary effect rather than proving synergy [56]. Demonstrating a true “1 + 1 > 2” interaction requires a dose-response matrix containing several concentrations of both the small molecule and the VHH or scFv. The resulting interaction may then be evaluated using Bliss independence, Loewe additivity, highest single agent, or zero interaction potency models. A primary model should ideally be specified in advance, with at least one additional model used for cross-validation. The term “synergy” should be reserved for combinations that consistently outperform the additive effect predicted by the selected reference models. Before entering animal studies, each combination should also meet several basic criteria. The small molecule should regulate its target and EV-related phenotype within a non-cytotoxic range. The VHH or scFv should be reproducibly produced and should specifically recognise native EV-associated antigens. The combined treatment should improve at least one immune-cell function without directly harming the recipient cells. Early animal studies may then focus on tolerability, pharmacokinetics, biodistribution, circulating EV markers, tumour growth, and immune-cell composition rather than immediately pursuing definitive therapeutic endpoints. Because CD81 and MET are also expressed in normal tissues, off-target tissue binding and systemic exposure require careful evaluation. CD81-directed binders, in particular, may be more suitable for ex vivo EV capture or enrichment until their tissue selectivity and safety are better understood. The purpose of the framework is therefore not simply to identify the apparently “strongest” molecule, but to determine, step by step, whether each intervention is biologically credible, experimentally reproducible, and worth advancing. Following experimental validation and identification of the authentic binding epitopes and critical contact residues, minimal binding motifs could be extracted and used to design antibody mimetics and functional peptides, or D-configured peptidomimetics, which, when combined with multivalent or bispecific display strategies [57], may yield EV-targeting binders that integrate antibody-like specificity, miniprotein-like stability, and peptidomimetic-like manufacturability.

A further contribution of this study is the development of an integrated and partially scriptable screening strategy for computationally generated VHH and scFv candidates. Recent advances in protein structure prediction, generative protein design, and sequence optimisation have greatly expanded the ability to explore antibody-like binders computationally. However, generative methods often produce large numbers of candidate structures, shifting the practical challenge from generating molecules to deciding which ones should be tested first. The present workflow integrates interface-contact analysis, hotspot recovery, structural-quality assessment, steric-clash detection, and developability-related features to reduce this candidate space. It can help direct limited experimental resources towards candidates with more coherent computational profiles and a stronger basis for further validation.

Overall, this study provides an extensible computational framework for exploring PDAC EV-associated immune regulation and narrows a broad candidate space to a more manageable set for future experimental work.

## Data and model availability statement

The dataset and candidate binder models generated in this study are available in the GitHub repository: https://github.com/ZhuYaojun1/PDAC-Qinba-exosome-dual-layer-intervention

## Declaration of interest

The authors declare no competing interests.

## Financial support statement

We gratefully acknowledge support from the Scientific Research Program of the Education Department of Shaanxi Province (23JK0359), the Natural Science Foundation of Shaanxi Province (2024JC-YBQN-0180).

## Authors’ contributions

Yaojun Zhu: Conceptualization, Methodology, Investigation, Formal analysis, Visualisation, Writing – original draft, Writing – review & editing. Xiaozhou Yang: Conceptualization, Writing – review & editing. Murtala Bindawa Isah: Writing – review & editing. Xiaoying Zhang: Conceptualization, Supervision, Funding acquisition, Project administration, Writing – review & editing. Yaojun Zhu and Xiaozhou Yang contributed equally to this work.

## Notes

### Competing Interest Statement

The authors have declared no competing interest.

